# Morphogen Patterning in Dynamic Tissues

**DOI:** 10.1101/2025.01.04.631293

**Authors:** Alex M. Plum, Mattia Serra

## Abstract

Embryogenesis integrates morphogenesis—coordinated cell movements—with morphogen patterning and cell differentiation. While largely studied independently, morphogenesis and patterning often unfold simultaneously in early embryos. Yet, how cell movements affect patterning remains unclear, as most pattern formation models assume static tissues. We address this gap by developing a mathematical framework for morphogen patterning in dynamic tissues, reformulating advection-reaction-diffusion models in cells’ reference frames—the most natural for signal interpretation and fate decisions. This framework (i) elucidates how morphogenesis mediates morphogen transport and compartmentalization: multicellular attractors enhance cell-cell diffusive transport, while repellers act as barriers, affecting cell fate induction and bifurcations. (ii) It formalizes cell-cell signaling ranges in dynamic tissues, deconfounding morphogenetic movements and identifying which cells can communicate. (iii) It provides two nondimensional numbers—typically distinct from the Péclet number—to assess when and where morphogenesis is relevant for patterning. (iv) It elucidates the generative role of cell density dynamics in patterning. We apply this framework to classic patterning models, morphogenetic motifs, and avian gastrulation data. Broadly, our work provides a quantitative perspective to rationalize dynamic tissue patterning in natural and synthetic embryos.

## Introduction

Embryogenesis requires the development of the embryo’s form–morphogenesis–and the diversification of its cells’ fates– differentiation [1]. In morphogenesis, tissues grow, stretch and flow, guided by molecular patterns [2,3] through which cells communicate and coordinate fate decisions. While morphogenesis and patterning are mostly studied separately, they often unfold simultaneously, especially in early embryogenesis, when morphogenetic movements are substantial and key patterns are established [1]. Cells commonly communicate via morphogens—signaling molecules transported between cells via diffusion or direct cell-cell transport. However, because morphogenetic movements rearrange tissue patches as signals travel through them, morphogenesis plays a confounding as well as generative role in patterning which is hard to disentangle [4–9]. While there are increasing studies on how the mechanics of morphogenesis influences cell fates [10–13], the effects of motion itself—the kinematics of morphogenesis—on cell fate coordination remains unclear [5].

The primary paradigms for patterning were developed for static tissues. Reaction-diffusion (RD) models [2, 14] rely on nonlinear interactions and differential diffusion to generate morphogen patterns that break symmetries or amplify weak asymmetries [15–19]. Positional information (PI), instead, refers to morphogen gradients cues cells use to infer their relative positions and generate discrete fate patterns [20]. RD and PI provide complementary perspectives on morphogen pattern formation and interpretation [21], typically assuming static tissues. Tissue growth affects RD dynamics [18, 19, 22] and PI [23–26]. However, embryogenesis involves complex tissue flows. Despite mounting evidence pointing to dynamic tissue patterning in embryogenesis [5, 6, 27–29], general mathematical frameworks are missing. Due to imaging advances, kinematic cell motion data (tissue velocities or cell trajectories) and patterning data (snapshots of gene expression, transcription factors, or signaling activity across tissues) are becoming increasingly available. An outstanding challenge is integrating noisy data on movement and molecules [29] to make sense of the feedback between morphogenesis and patterning. Even with adequate data, the core conceptual question of dynamic tissue patterning remains: how does morphogenesis affect the signals cells receive? Morphogens activate intracellular feedback so that cell fates are affected by their morphogen exposure over time [30–32]. The combination of motion and memory calls for patterning frameworks describing morphogen exposure along cell trajectories.

Theoretically, this requires a nonlinear reference frame change from standard fixed Eulerian coordinates (**x**, Fig. 1A) to Lagrangian coordinates—co-moving with cells (**x**_0_ or **x**_*t*_, Fig. 1B). Throughout, we bold vectors and matrices, but not scalars. Unlike Eulerian coordinates, Lagrangian coordinates are always associated with the same cells, even as they move. Fig. 1C depicts a morphogen concentration 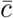—overbar indicates functions of Eulerian coordinates **x**—at different times (black curves). As cells (red, white, and blue) move through 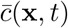, they experience a dynamic morphogen exposure *c*(**x**_0_, *t*). The confounding effects of morphogenesis on patterning arise because 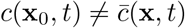, and fate decisions happen in the cell’s reference frame—i.e. in Lagrangian coordinates—taking *c*(**x**_0_, *t*) as input (Fig. 1D). In addition to affecting the positions sampled over time, morphogenesis may affect the morphogen concentrations at those positions by reshaping morphogen gradients. It is not obvious how the interplay of diffusion and deformation (the relative motion of different tissue patches) affects *c*(**x**_0_, *t*) (Fig. 1E). Here, we develop a mathematical framework for the dynamics of *c*(**x**_0_, *t*), revealing intimate connections on how morphogenesis mediates morphogen transport and exposure.

**Figure 1:**
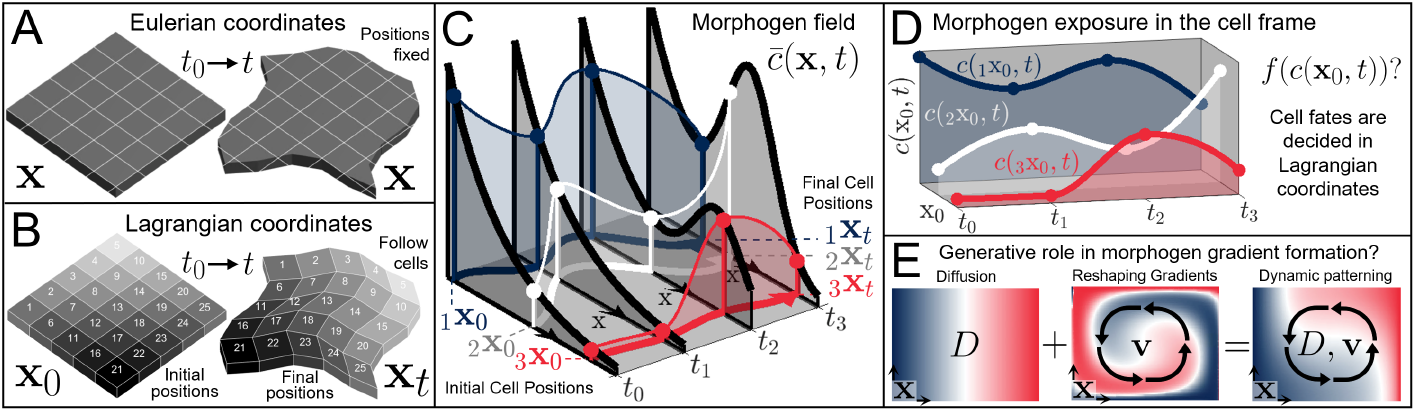
Dynamic Tissue Patterning. A) Eulerian coordinates represent fixed positions in space through which cells and morphogens move. B) Lagrangian (cell) coordinates co-move with cells and are associated with labeled tissue patches over time. C) 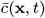 (black curves) show a morphogen concentration in Eulerian coordinates. *c*(**x**_0_, *t*) (colored curves) mark morphogen exposure of three cells moving from _*i*_**x**_0_ to _*i*_**x**_*t*_ (Movie 1). D) Gene expression dynamics and cell fate decisions depend on complicated functions *f* (*c*(**x**_0_, *t*)) of morphogen exposure over time. E) Even in simple models where fates follow directly from morphogen patterns (e.g., the fixed gradient solution to the French-Flag problem [20]), morphogenesis can affect fate patterns (in **x**) by reshaping morphogen gradients. Left: Diffusion from a source on the left boundary of a 2D epithelium form a morphogen gradient colored with high (blue), medium (white), and low (red) morphogen concentrations. Middle: Tissue motion (morphogenesis, black arrows) rearranges this pattern. Right: The combined effect of diffusion and tissue motion generates a different pattern, distinct from mere rearrangement or static gradient formation.

## Results

In static patterning, 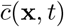 is described by the diffusion equation 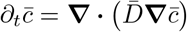 with isotropic diffusivity 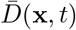, typically complemented by source, degradation, and reaction terms. When advective motion is present, 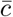 obeys the advection-diffusion equation

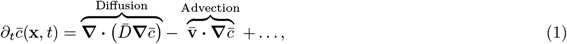

where 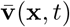 is an incompressible 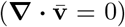 velocity representing morphogenetic movements. Whenever possible, we omit the explicit (**x**, *t*) dependence. Throughout, we assume that morphogens co-move with cells due, for example, to extracellular matrix (ECM) co-movement, common in early amniotes [33]. Later, we incorporate the effects of compressibility, cell divisions and deaths, and relative tissue-ECM velocities (*SI Sec. 1*).

### Morphogen exposure in the cell frame

We rewrite Eq. (1), changing coordinates from Eulerian **x** to Lagrangian **x**_0_, using the coordinate change 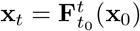 corresponding to cell trajectories (Fig. 1A-B). Because Lagrangian coordinates map between cell initial positions **x**_0_ and later positions **x**_*t*_, one can visualize patterns on the initial (undeformed) or final (deformed) configurations. This resonates with “fate mapping” techniques in classical embryology in which initial embryonic regions are labeled according to where they go. Equation (1) for 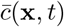 transforms (*SI Sec. 1*) [34–36] into Eq. (2) for *c*(**x**_0_, *t*), the time-*t* morphogen exposure of a moving tissue patch that started at (**x**_0_, *t*_0_):

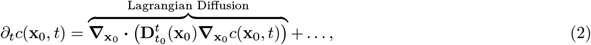

where **∇**_x0_ denotes derivatives with respect to **x**_0_. In Lagrangian coordinates, the Eulerian advection and diffusion terms reduce to a single term with an equivalent Lagrangian diffusion tensor

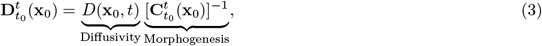

where 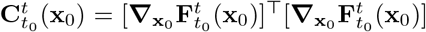 denotes the Cauchy-Green strain tensor, a symmetric positive definite (i.e. with real orthogonal eigenvectors) positive definite (i.e. with real positive eigenvalues) matrix, and 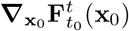 is the deformation gradient tensor. Therefore, in the cell frame, cell-cell morphogen transport blends the Eulerian diffusivity sampled along trajectories 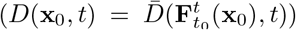 with the effect of their deforming environment, encoded in 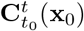. There is no advection term in the cell frame, absorbed in the material time derivative *d*_*t*_*c*(**x**_0_, *t*) = ∂_*t*_*c*(**x**_0_, *t*). However, the spatial coordinate change **x** → **x**_0_ has a nontrivial effect on the non-cell-autonomous term **∇** · (…) (Eq. (1), *SI Sec. 1*). Strikingly, Eq. (3) reveals that cell-cell transport in the cell frame inherits *i*) the space dependence, *ii*) anisotropy (direction dependence) and *iii*) strength of deformation of its co-moving environment. Without deformation, 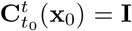, recovering isotropic Eulerian diffusion 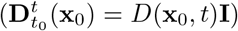 or 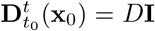 if *D* is uniform in space. Note that in Eq. (1) and Eq. (2) the effective molecular diffusivity *D* may be locally regulated along trajectories by local factors such as cell number density [37] or ECM component concentrations [10]. Hereafter, we consider a fixed *D* without loss of generality and note that our framework applies to heterogeneous *D*.

### Connection to the Dynamic Morphoskeleton

The relationship between 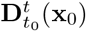 and 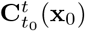 provides an exact, profound connection between dynamic tissue patterning and the Dynamic Morphoskeleton (DM) [38, 39], a frame-invariant, compressed representation of complex tissue flow that has been applied across model systems in developmental biology and active matter [38, 40–46]. The DM reduces noisy cell trajectory data to a robust set of tissue flow organizers: attracting and repelling coherent structures [47], hereafter just *attractors* and *repellers*. The largest eigenvalue 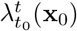 of 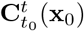 quantifies the maximum separation of cells starting near **x**_0_ along the direction of its corresponding eigenvector 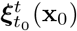. High values of 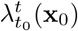 locate repellers, i.e. initial (**x**_0_) tissue regions where nearby cells will maximally separate by time *t* (Fig. 2A-B). Conversely, performing the analysis in backward time, high values of 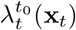 mark attractors, i.e. final (**x**_*t*_) locations towards which initially separated cells will converge by time *t* (Fig. 2A-B) [38]. The DM can be extracted for tissue flows in 1, 2, or 3 dimensions. In 1D flows, attractors and repellers are dynamic points (Fig. 2B). In 2D flows, our focus here, attractors and repellers are dynamic 1D curves (Fig. 2A and Fig. 4). Arising from the cumulative deformation along trajectories, the DM is undetectable from inspection of 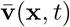, even in 1D (Fig. 2B). See SFig. 3 for a connection between Eulerian analysis and the DM using the experimental flow data from Fig. 4.

**Figure 2:**
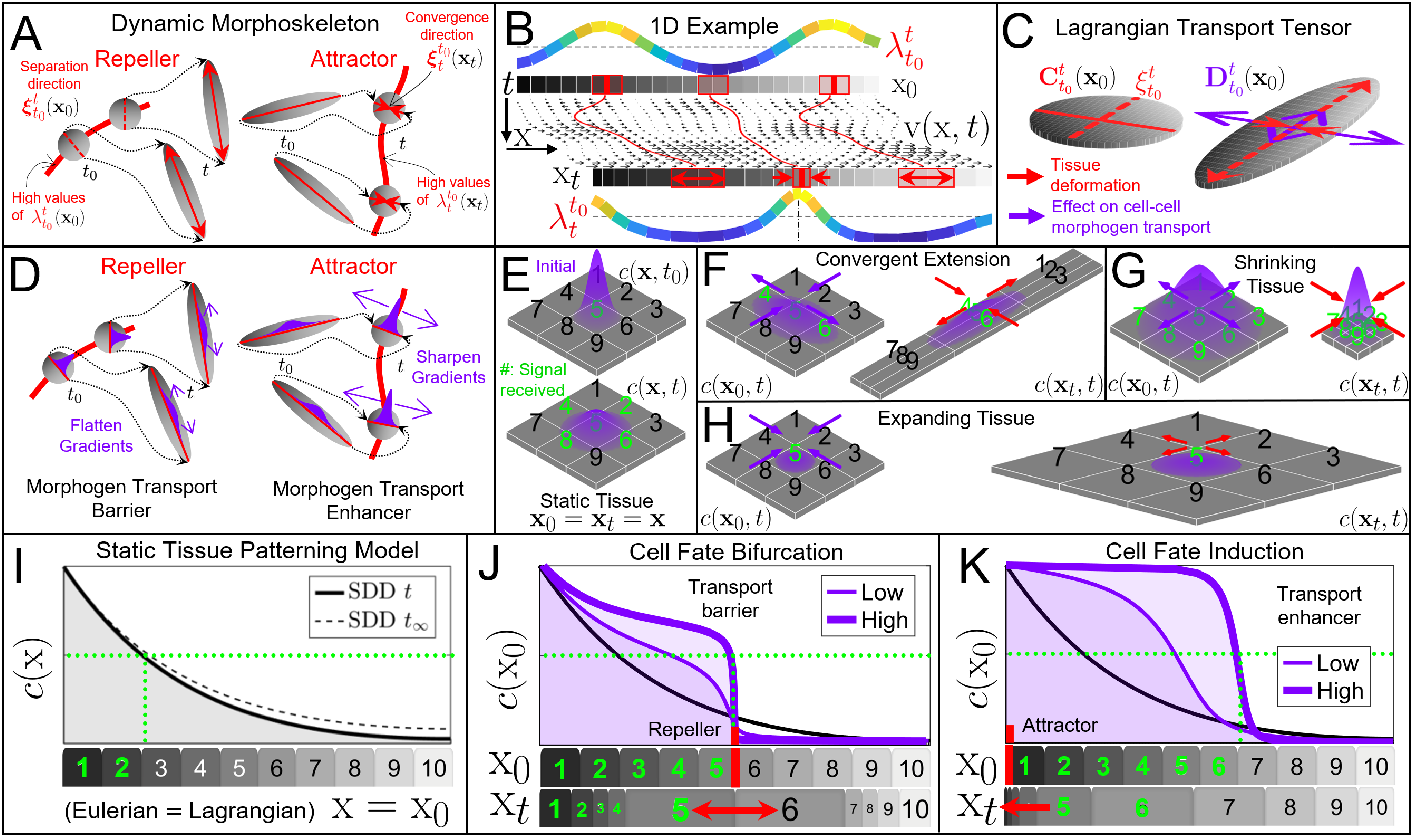
The Dynamic Morphoskeleton mediates intercellular morphogen transport in the cell frame. A) DM consists of attractors and repellers. High values (ridges) of 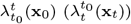 mark repellers (attractors) across which cells will maximally separate (converge) along the direction 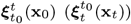 during the time interval [*t*_0_, *t*]. Repellers (attractors) are visualized at the initial **x**_0_ (final **x**_*t*_) tissue configurations. *λ*, ***ξ*** denote the largest eigenvalue and eigenvector of the Cauchy–Green strain tensor **C**. B) DM for simulated, 1D time-dependent velocity data consisting of two repellers and one attractor (red). C) Visualization of 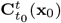 and 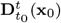 (Eq. (3)) for the linear convergent extension flow (Eq. (4)). In the Lagrangian frame, diffusive transport is enhanced (diminished) in the contracting (extending) direction. D) Repellers (attractors) reduce (enhance) diffusive transport across them. E-H) Visualization of diffusive morphogen transport in the cell frame for static and uniformly deforming tissues. E) An initial Gaussian of morphogen concentration *c*(**x**, *t*_0_) with max concentration *c*_max_ and variance 0.01 diffuses (*D* = 0.01, no flux boundary conditions) above a static 2D epithelium (length 1), simulated from *t*_0_ = 0 to *t* = 1 (all arbitrary units). Tissue patch labels turn green if average *c*(**x**_0_, *t*) > *c*_max_/50. F) Simultaneous diffusion and convergent extension (**v** = [−x, y]) with the same initial condition and parameters as E. Final morphogen concentrations depicted in Lagrangian coordinates at initial positions (*c*(**x**_0_, *t*), left) and final positions (*c*(**x**_*t*_, *t*), right). Tissue deformation reshapes morphogen gradients, mediating cell-cell diffusive fluxes in the cell frame. G, H) Same as F for a uniformly shrinking (**v** = [− x, y]) and expanding (**v** = [x, y]) tissues. Movie 2 shows the time evolution of panels E-H. I) Synthesis, diffusion, and degradation model 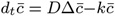. Starting at *t*_0_, morphogens are secreted at the left boundary of a 1D domain with constant morphogen flux 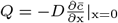, and no flux at the right boundary. Within the domain, morphogens diffuse and degrade uniformly (rate *k*). The solid black curve shows concentrations at *t* = *k*^−1^ with a fixed source at the left boundary and target tissue patches (1-10), colored green if concentrations exceed a morphogen threshold (green dotted line). The dashed curve is the steady-state solution 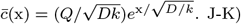 Same as I with the addition of steady, nonuniform flows with low or high deformation rates generating a strong repeller (J) and attractor (K), marked in red. Final deformed configurations x_*t*_ are shown below for high deformation rates. For the time evolution, see Movie 3. J) The repeller limits diffusive morphogen transport across it. K) The attractor enhances diffusive morphogen transport away from it. Parameters: I-K) uniform *D* = 0.001, uniform *k* = 0.01, *t* = 100, *L* = 1. Fixed *c*(0) on the left boundary (arbitrary units) and no flux on the right boundary. See SFig. 4A for lower deformation rates and velocity fields used in J-K.

To interpret the connection between deformation 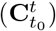 and 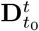 (Eq. (3)), consider a linear, time-independent convergent extension velocity field 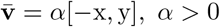. Near the origin, the x-axis (y-axis) represents a repeller (attractor) as trajectories on opposite sides will separate (converge) (Fig. 2C). For this flow, 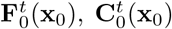 its eigenvalues 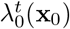 (largest) and 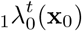 (smallest), and 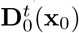 are:

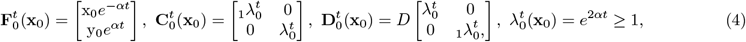

where 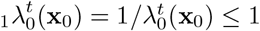 due to incompressibility. It is instructive to compare the eigenvalues and (orthogonal) eigenvectors of 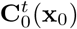 and 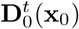. The largest (smallest) eigenvector 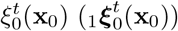 of 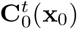 coincides with the smallest (largest) eigenvector of 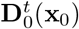, implying that the local contracting direction experiences enhanced cell-cell transport 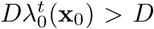, while the expanding direction experiences reduced cell-cell transport 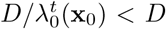 (Fig.2C). These effects were recently experimentally realized, imprinting directionality on chemical waves [48]. The same conclusions hold for general nonlinear and time-dependent flows, where regions of high 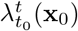 define the DM (Fig. 2A), and the connection between 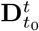 and 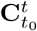 (Eq. (2)) reveals that repellers (attractors) act as barriers (enhancers) to morphogen transport from cell to cell (Fig. 2D). These connections have been recognized in fluid mixing research [34–36], but not in the biological context.

It is key to recognize that *i*) 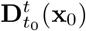 parametrizes only non-cell-autonomous mechanisms, distinct from the advective transport of *c*: for *D* = 0, no morphogens are transported between cells. *ii*) The modulation of morphogen transport in the cell frame is due to tissue deformation, leaving the molecular diffusivity *D* unaltered. To visualize this effect, we solve Eq. (2) using the linear convergent-extension flow (Eq. (4)), an initial centered Gaussian *c*(**x**_0_, *t*_0_) (Fig. 2E top). Without deformation, 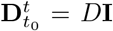 and *c*(**x**_0_, *t*) spreads isotropically (Fig. 2E bottom), reaching (i.e. *c* > *c*_threshold_) four neighboring tissue patches (green) at time *t*. With deformation (*α* > 0), 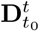 becomes anisotropic (Eq. (4)), enhancing (diminishing) morphogen transport across the attractor (repeller) (Fig. 2F). In the cell frame at the initial time (**x**_0_), *c* reaches more cells along the direction of convergence than of extension (Fig. 2F, left). This non-cell-autonomous effect arises because tissue deformation sharpens and flattens morphogen gradients, modulating the diffusive fluxes between cells.

Visualizing morphogen exposure and transport in the cell frame (**x**_0_, 2F left) de-confounds the effects of tissue flows, which become overwhelming for general flows (see Fig. 4 and Movie 4). Crucially, the effects of morphogenesis on cell-cell transport are not specific to this simple convergent-extension flow (Eq. (4)). In general, time-dependent nonlinear flows with arbitrary morphogen distributions and additional cell-autonomous terms for sources, sinks, and reactions, these same insights about barriers and enhancers to diffusive cell-cell morphogen transport remain encoded in 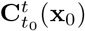 (Eq. (3)), which quantifies morphogenesis and can be computed from experimental data [38, 42, 45].

### Isotropic vs. anisotropic deformation

In incompressible flows, deformation is purely anisotropic (Fig. 2C). However, for compressible 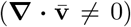 flows (e.g., due to cell division, death and expansion), deformation have anisotropic and isotropic components encoded in 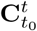, both affecting 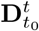. We omit the dependence on **x**_0_ whenever possible. In general, 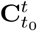 has two eigenvalues, 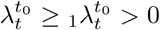. To isolate isotropic and anisotropic effects, we use spectral decomposition and diagonalize 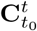 in its eigenbasis (*SI Sec. 2*)

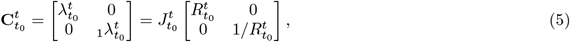

where 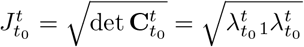 and 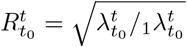. To first approximation, a small undeformed circular tissue patch is deformed by tissue flows into an ellipsoid (Fig. 2C and Fig S2). 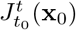 represents the ratio of final (*t*) to initial (*t*_0_) area of the tissue patch started at (**x**_0_, *t*_0_), while 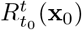 denotes the ratio of ellipse major (along 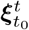) and minor axes (perpendicular to 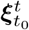), quantifying anisotropic deformation (*SI Sec. 2*). In incompressible flows (Eq. (4), Fig. 2F), 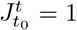. Conversely, in purely isotropic deformation 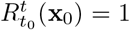: i.e. 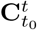 does not have distinct eigenvalues and eigenvectors. General tissue flows produce both except at distinct locations where deformation is isotropic: 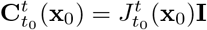 (SFig. 2).

### Patterning in compressible tissue flows

For compressible flows, Eq. (1) gains an additional term 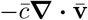. In Lagrangian coordinates, Eq. (2) then becomes

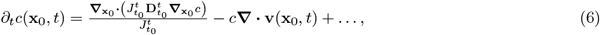

where 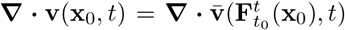 indicates the velocity divergence sampled across trajectories. Eq. (6) shows that also in compressible flows, the Lagrangian diffusion tensor inherits the anisotropy of its deforming environment, and 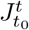 provides an additional isotropic modulation of cell-cell transport magnitudes (*SI Sec. 1*). To gain intuition, consider the compressible linear convergent flow **v** = −*α*[x, y], *α* > 0, for which 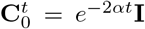 and Eq. (6) gives 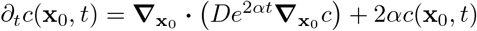. Isotropic shrinkage enhances diffusive fluxes, spreading signals to more cells (Fig. 2G). Conversely, isotropic expansion (*α* < 0) confines signals to fewer cells (Fig. 2H).

### Dynamic cell fate bifurcations and fate induction

In development, morphogens are synthesized and degrade. In the classic synthesis, diffusion, and degradation (SDD) model, morphogens are locally produced at one boundary and diffuse and degrade (rate *k*) uniformly on a static domain [49, 50]. In Fig. 2I, the dashed black curve marks the steady state solution 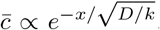, while the solid black curve shows a finite-time numerical solution 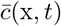.

In dynamic tissue patterning, source and target cells may not maintain a fixed distance. However, even in the simplest case, when they do, deformation between them may affect their communication. To this end, we add to the SDD model a stationary compressible flow with a repeller between the source (left boundary) and target cells on the right (red segment in Fig. 2J) but zero velocity at the boundaries so that the spatial domain remains fixed. We display the solution *c*(**x**_0_, *t*) (purple curve in Fig. 2J) for different strengths of the repeller (i.e. deformation rates). The repeller acts as a diffusion barrier, creating a steep gradient of *c*(**x**_0_, *t*): at time *t*, cells initially sitting on either side of the repeller will be exposed to markedly different morphogen concentrations. The black curve is as in Fig. 2J. This simple example illustrates repellers’ capacity to aid in morphogen compartment formation due to a non-cell autonomous effect caused by tissue deformation, contrasted with mere advection. On the right of the repeller, for example, the velocity (advective transport) points to the right. Yet, cells on the right of the domain experience reduced exposure to morphogens from source cells. These findings resonate with recent experimental evidence associating repellers with cell-fate bifurcations during zebrafish body axis elongation at 12 hpf [42] and in chick embryogenesis at HH4-HH8 [44].

Another dynamic tissue patterning motif is the continual induction of cell fates as cells move through dynamically stable morphological structures [51]. For example, in avian gastrulation, cells move towards and ingress through the primitive streak (PS), a sharp attractor [38]. As cells internalize, they complete epithelial-to-mesenchymal transitions (EMT) that begin before they reach the attractor and involve non-cell-autonomous induction [51]. At its simplest, this process can be described using the SDD model with advective terms and a morphogen source (mesoderm cells pre-internalization) co-located with a sharp attractor on the left boundary (Fig. 2K). Figure 2K displays the solution *c*(**x**_0_, *t* = *k*^−1^) for different deformation rates, showing how the attractor enhances transport of *c*(**x**_0_, *t*) from the source, resulting in a steep rightward-expanding front. The left of the front, where *c* is high, labels induced cells. The advection is to the left (see **x**_*t*_), yet the attractor enhances cell-cell diffusive transport to the right.

### Criteria for morphogenesis-mediated tissue patterning

Our framework relies on the following assumptions: First, morphogens diffuse through the extracellular matrix (ECM) near cellular surfaces [52], limiting free diffusion away from the tissue. This ubiquitous role of the ECM [10] may explain why adding ECM components (e.g. Matrigel) can be crucial for complex patterning in synthetic embryo models [7]. Second, the ECM moves with the underlying tissue, as demonstrated experimentally in the gastrulating avian embryo [4] and in *Xenopus* [53]. When cells and ECM are not co-moving [54,55], our framework can be adapted by accounting for their relative velocity (*SI Sec. 1*). While we focus on extracellular diffusion, our framework applies to any transport mechanisms modeled by diffusive fluxes (Eq. (1)).

Next, we provide two conditions to assess if tissue flows may impact patterning during a time interval of interest [*t*_0_, *t* = *t*_0_ + *T*]: *i*) Morphogenetic movements must generate appreciable deformation, causing deviations from simple diffusion in the cell frame: 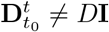, which requires 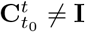 (Eq. (3)). *ii*) Diffusion must not quickly flatten gradients over the region of interest before deformation effects manifest. The first condition can be expressed as Λ^−1^ < *T* where 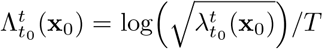 is the finite-time Lyapunov exponent (FTLE), evaluated where deformation is highest, i.e. at repellers or attractor (*SI Sec. 3 and SI Appendix 1*). Λ^−1^ is the Lyapunov time, which indicates the time taken for a small initial undeformed tissue patch at **x**_0_ to undergo appreciable deformation—i.e., Λ^−1^ represents a deformation time scale (*SI Sec. 3*). In simple linear flows like Eq. (4), Λ = *α*, and Λ^−1^ < *T* ⟹ *α*^−1^ < *T*, meaning that for short *T*, deformation rates must be stronger (larger *α*) to achieve 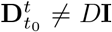. Alternatively, for a given deformation rate *α*, one needs *T* > *α*^−1^ to observe 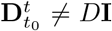 [56, 57]. For nonlinear stationary or non-stationary flows, Λ is the effective *α* and can be computed from experimental tissue velocities [38] or cell trajectories [39] for any time interval of interest. SFig. 4 shows the effect of varying Ω_1_ associated with Fig. 2J-K and Ω_1_ from avian gastrulation data (Fig. 4).

If patterning mechanisms sustain nonzero concentration gradients, as in the SDD model (Fig. 2I), then *ii*) is automatically satisfied, and *i*) is the only condition that needs to be checked. By contrast, if no mechanisms are known to sustain ∇*c* ≠ 0, one needs that the deformation time scale Λ^−1^ is smaller than the diffusion time scale *L*^2^/*D* [3], where *L* is the characteristic length of the region of interest for patterning (*SI Sec. 3*). We summarize these conditions as

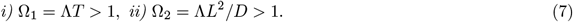

Ω_1_ and Ω_2_ are frame invariant, non-dimensional parameters, and the latter differs from the classic Péclet number (Pe = |**v**|*L*/*D*), a ratio of advective and diffusive transport rates, which does not account for deformation. In the Lagrangian frame (Eq. (2)), advection is absent, and a uniform velocity (Pe > 0) generates no deformation (Ω_2_ = 0). Ω_2_ can coincide with the alternative Pe = *αL*^2^/*D* based on strain rates (*α*), but only in two specific cases: for linear stationary flows as in Eq. (4) or for general flows over infinitesimally short time intervals [*t*_0_, *t* = *t*_0_ + δ*t*] (*SI Sec. 3*). By contrast, Ω_2_ applies to general, nonlinear morphogenetic processes over finite times. It is key to recognize that contrary to Eulerian advection-reaction-diffusion equations (Eq. (1)), morphogen dynamics in the moving cell frame depend on tissue deformation (Eq. (6)), leading to nondimensional parameters that encode cumulative deformation instead of instantaneous motion or stretching rates. For an algorithmic procedure to verify Eq. (7) from experimental data, see *SI Algorithm 1*. Finally, if the fastest mechanism to eliminate gradients is not diffusion, *L*^2^/*D* in Ω_2_ should be replaced by the relevant time scale, e.g., degradation *k*^−1^. Alternatively, if a relay mechanism [58–60] drives morphogen spreading, it should be replaced by *L*/*w*, where *w* is the effective front speed. Conditions (7) are typically satisfied in early embryogenesis (see avian gastrulation below and SFig. 5).

### Embryological Light Cones

The above results demonstrate how morphogenesis modulates diffusive fluxes (Fig. 2C-H) and can thereby influence which cells communicate (Fig. 2J-K). To formalize this effect, we ask: If a cell 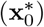 sends signals via morphogen transport over [*t*_0_, *t*_0_ + *T*], which cells could receive them? Similarly, from which cells could 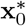 have received signals sent during a past time interval [*t*_0_ −*T, t*_0_]. We define these regions over increasing durations *T* as “Embryological Light Cones” or 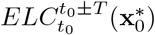. In cosmology, light cones bound domains of causal interaction and have been adapted in pattern formation in automata models [61, 62]. Whereas classic light cones invoke a constant speed, the rate of molecular information propagation from cell to cell in dynamic tissues varies, modulated by morphogenetic movements 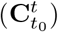. Accordingly, ELCs can be curvilinear instead of conic but retain apices at 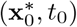 and expand into the past and future. Future cones can be estimated by evolving *c*(**x**_0_, *t*), considering a source of *c* at 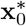 and subject to known or candidate signal propagation mechanisms over [*t*_0_, *t*]. The ELC bounds the set of cells (**x**_0_, *t* > *t*_0_) exposed to morphogen concentrations above a threshold *c*_min_ (or threshold gradients etc.) based on thresholded cellular responses. Similarly, estimating past cones requires determining retrospectively the set of cells (**x**_0_, *t* < *t*_0_) for which 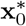 lies in their future cones at *t*_0_.

In static tissue patterning, ELCs expand isotropically with time-dependent ELC radius *r* and are time-symmetric. Figure 3A shows ELCs for diffusion 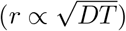; a production relay (*r* ∝ *wT*); and the SDD model, where the ELC expands to a fixed radius 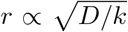 (*SI Sec. 4*), formalizing the classical notion of morphogen range [63, 64]. In dynamic tissue patterning, signaling mechanisms rely on cell-autonomous sources, sinks and reactions, augmented by non-cell-autonomous communication mechanisms modeled by Eqs. (2,6), which affect ELCs. ELCs are no longer time-symmetric, even with constant deformation rates (Fig. 3B, *SI Sec. 4*), because continuous deformation introduces an explicit-time dependence to transport rates (consider for example Eq. (4)).

**Figure 3:**
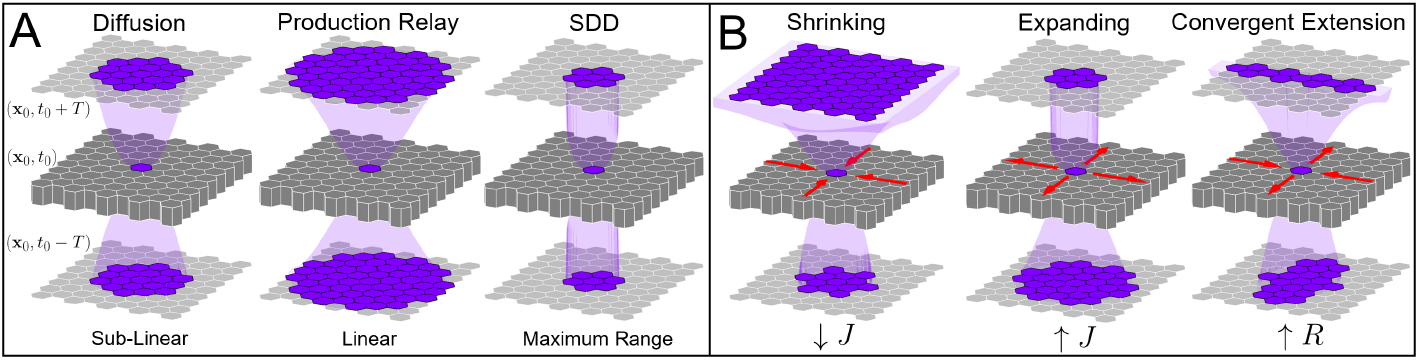
Embryological Light Cones reveal cell-cell interaction ranges. Past and future cones for tissue patches at 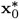, *t*_0_ depend on known or candidate signaling mechanisms and tissue deformation. A patch’s future 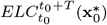 (top) delimits cells (**x**_0_) at *t* = *t*_0_ +*T* that can receive morphogen signals from cells at 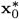. A cell’s past cone 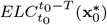 (bottom) delimits cells that could have sent signals to (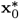, *t*_0_) from (**x**_0_, *t*_0_ − *T*). ELCs simulated with a fixed source concentration *c* = 1 and threshold *c*_min_ = 0.1 in different morphogen transport scenarios. A) ELCs in static tissue patterning with diffusion (*D*), a production relay (front speed *w* ≈ 0.8, *SI Sec. 4*), and in the SDD model (degradation rate *k* = 2). B) ELCs in dynamic tissue patterning with diffusion and stationary, isotropic negative (**v** = −0.8[x, y]), positive (**v** = 0.8[x, y]), and anisotropic **v** = 0.8[x, −y] deformation rates. All 2D simulations use parameters: *D* = 0.2, *t*_0_ = 0, *T* = 1, and *L* = 2 (tissue size). For analytical predictions, see *SI Sec. 4-5*.

In an isotopically shrinking tissue (*J* < 1), future cones can grow super-superlinearly, enhancing cooperativity, while past cones are narrower. The opposite occurs in an isotropically growing tissue (*J* > 1), where past cones can expand super-linearly while future cone expansion slows. This reflects cells’ waning global influence within a growing tissue and the efficiency of patterning before extensive growth [1]. When deformation is anisotropic (*R* > 1), past and future cones expand anisotropically, with enhanced and diminished ranges determined by the directions of convergence and extension (Figs. 2C,F). Next, we compute ELCs using highly remodeling avian gastrulation flows.

### Avian gastrulation example

Avian gastrulation involves fast, compressible, time-dependent tissue flows **v**(**x**, *t*) over 12*h* (Fig. 4A) that change the embryo’s shape and form the primitive streak (PS) along the embryo’s anterior-posterior (A-P) axis. During gastrulation, cells signal via morphogens such as Nodal, Vg1, BMP, Wnt, and FGF [66], which may be transported directly via diffusion through the sub-epiblastic ECM, co-moving with cells [4], or be limited to shorter-range paracrine signaling [67]. To elucidate the role of morphogenesis in morphogen transport, we consider *t*_0_ = 0*h* (developmental stage HH1) and *T* = 12*h* (stage HH3). Next, we compute the DM from **v**(**x**, *t*), consisting of two repellers (Fig. 4B) and one attractor (Fig. 4C), marking regions of highest deformation quantified by 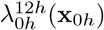 and 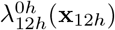 [38].Repeller 1 marks the initial position (**x**_0*h*_) of the embryonic-extraembryonic boundary. The PS is marked by the attractor at the final embryo configuration (**x**_12*h*_). It is also informative to visualize its domain of attraction: i.e., the initial cells (**x**_0*h*_) that will eventually converge to the attractor (within red dotted curve in Figs. 4B,D). Finally, Repeller 2 (Fig. 4B) splits the anterior-posterior domain of attraction that will converge to the anterior and posterior PS. Using Eq. (7), on the DM, Λ^−1^ ≈ 4*h*, hence Ω_1_ ≈ 3 (SFig. 4), revealing 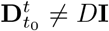 after *t* ≈ 4*h* and where these deviations are strongest. Typical morphogen diffusivities are *D* = 0.1 − 100 *µm*^2^ *s*^−1^ [52]. Assuming diffusion-limited transport with *D* = 1 *µm*^2^ *s*^−1^ and *L* ≥ 200*µm* (≈ 20 cell diameters), Ω_2_ > 3.

**Figure 4:**
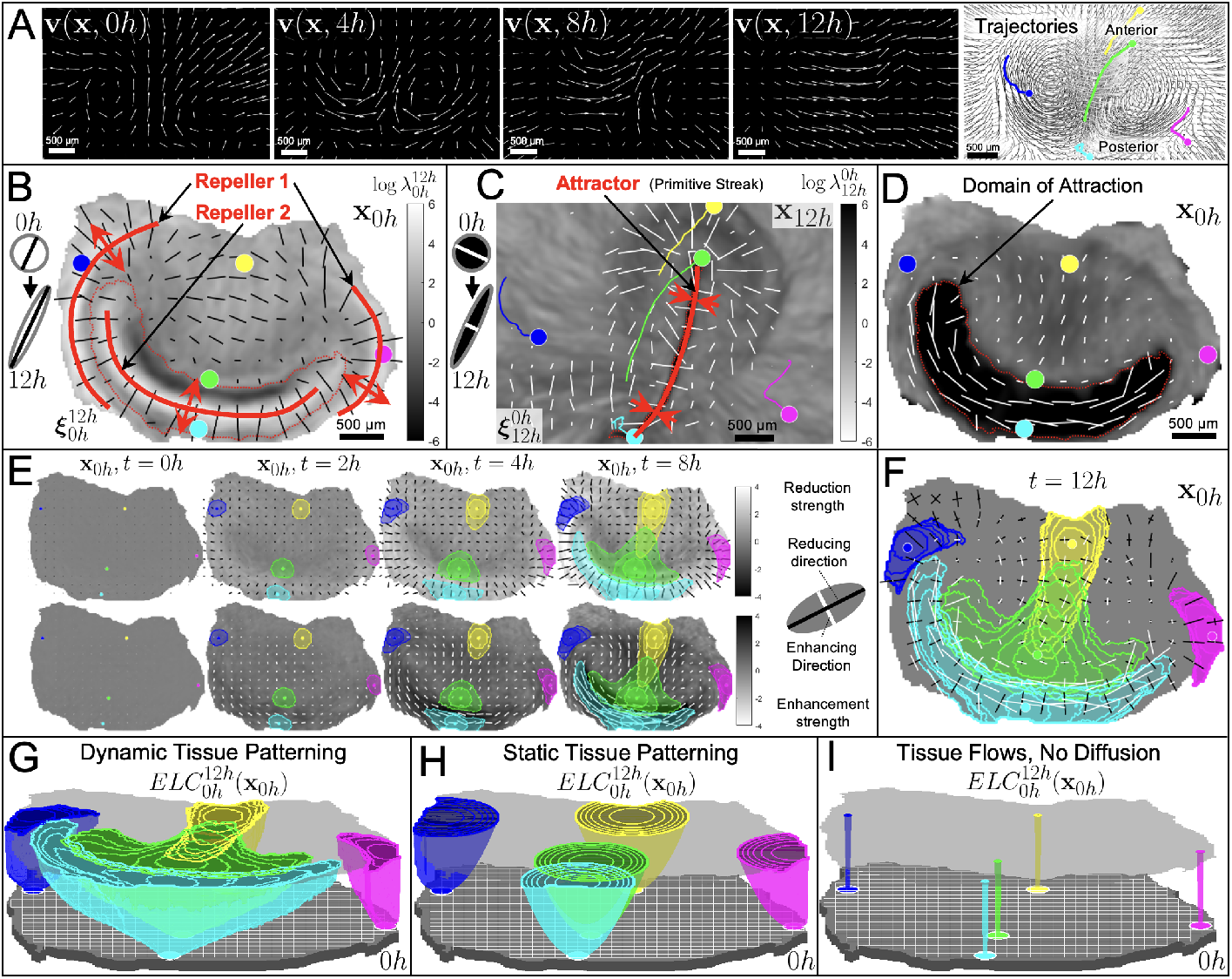
Morphogenesis-Mediated Morphogen Transport in Avian Gastrulation Flows. A) Tissue velocities in the gastrulating chick embryo (*t*_0_ = 0*h* corresponds to developmental stage HH1) and *t* = 12*h* (stage HH3), obtained via particle image velocimetry [65]. From **v**, we compute cell trajectories (right) and tissue deformation encoded in **C** [38]. Source trajectories are depicted in colors with markers at their final positions. B-D) Analysis over the full 12*h* reveals two repellers and one attractor (the Dynamic Morphoskeleton [38]), along with directions (**C** eigenvectors) of convergence (white) and separation (black) with lengths proportional to their corresponding eigenvalues. B) Two repellers (transport barriers) marked by high values of 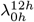 on **x**_0*h*_. Bars indicate the direction of maximum separation 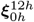 with length proportional to log 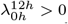. C) One attractor (transport enhancer) marked by high values of 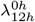 on **x**_12*h*_. Bars indicate the direction of maximum convergence 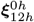 with length proportional to log 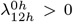. D) Same as C, displayed at the initial embryo configuration **x**_0_ (colorbar as in C). E-F) Simulations with sources secreting morphogens at rate *Q*, which diffuse with *D* = 1*µm*^2^/*s*. Morphogens and cells move with **v**(**x**, *t*) (A). E) Top (bottom) row depicts the maximally stretching (shrinking) directions as in B,D. Scalar fields on the top (bottom) depict the strength of cell-cell diffusion reduction (enhancement) on a log scale as in B,D. Colored contours mark tissue patches at their **x**_0*h*_ positions raised above a threshold morphogen concentration *Q*/100 at *t* = 0, 2, 4,and 8*h*. Note that these results do not depend on the value of *Q* > 0. F) Contours from E along with the outermost contour and diffusive transport reduction (enhancement) directions corresponding to *t* = 12*h*. White (black) bars mark the direction of diffusion enhancement (reduction) proportional to their length. G) Each source’s future 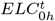 over increasing *t*. Nested contours correspond to ELCs at (*t* = 2, 4, 6, 8, 10, and 12*h*). Movie 5 shows the time evolution of cell trajectories and dynamic tissue patterning associated with B-G. H) Same as G for static tissue patterning (no cell motion: **v** = **0**) using the same parameters as in G. I) Same as G, where tissue flows are present but *D* = 0, highlighting that the role of morphogenesis in mediating non-cell autonomous (or cell-cell) signaling is not due to cell-autonomous morphogen rearrangements.

To elucidate the strength and anisotropy of deformation over [0*h*, 12*h*], we use the eigenvalues and eigenvectors of **C** (Eq. (5)), displayed both on **x**_0*h*_ and on the deformed embryo configuration **x**_12*h*_ (to relate quantities at **x**_0*h*_ and **x**_12*h*_, see SFig. 2). Fig. 4B shows log 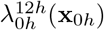, i.e. the strength of diffusion reduction (where > 0) along the stretching direction (black bars), highlighting that repellers reduce diffusive fluxes between separating cells. Fig. 4D conversely reveals the directions (white bars) along which tissue patches shrink the most, showing that there is an enhanced diffusive flux (colorbar as in Fig. 4C, where > 0) perpendicular to the A-P axis within the domain of attraction. It is instructive to visualize this information also at the final embryo configuration **x**_12*h*_. Fig. 4C shows the strength of diffusion enhancement (where log 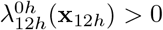) and the direction (white bars), highlighting how diffusive fluxes are mostly enhanced laterally at the attractor, and mildly along the radial direction in the fountain-shaped region surrounding the anterior part of the PS and emanating from the Henson’s Node. Because they exhibit the strongest modulation of cell-cell morphogen transport, repellers and attractors are the essential kinematic features that mediate cell-cell communication, reshaping ELCs.

To visualize our predictions, we examined tissue patches (colored disks in Figs. 4A-D) initialized *i*) outside Repeller 1 (blue and magenta) and therefore in the extraembryonic tissue, *ii*) in the anterior embryo (yellow), and *iii*) anterior and posterior of Repeller 2 (green and cyan). We treat these five patches as morphogen point sources, secreting morphogens at a fixed rate during [0*h* − 12*h*], using a typical morphogen diffusivity *D* = 1*µm*^2^/*s* (see SFig. 5 for alternative values). The source patches move with the tissue (Fig. 4A,C), but remain fixed in the cell frame (Fig. 4B,D). In essence, our framework casts the dynamic tissue patterning in gastrulation, typically challenging to rationalize due to the complex flows, as an equivalent static tissue patterning problem over the initial embryo configuration **x**_0*h*_, with the effects of cell motion on cell-cell signaling encoded in 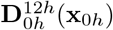. Fig. 4E shows the temporal evolution of the DM and 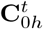 for increasing *t*, decomposed as in Figs. 4B,D. Overlaying the deformation fields, we plot a top-down view of each source’s future 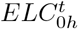, with colored contours marking the extents at each intermediate time. While initially expanding slowly in all directions 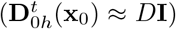, the effects of cumulative deformation manifest in the expansion of ELCs around 4*h*—when 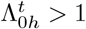 on the DM—and later become more dramatic. For example, before 4*h*, the green ELC expansion is mostly isotropic (expanding as a circle); after 4*h*, diffusive fluxes become enhanced across the posterior, especially perpendicular to the A-P axis (Fig. 4F).

Fig. 4F depicts the ELCs as colored contours for 2*h t* intervals (as in E) and the final outermost contour at 12*h* along with directions of maximum deformation (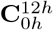 eigenvectors). White (black) bars mark directions of enhanced (reduced) diffusion with bar lengths indicating the strength of this modulation. This panel also reveals the degree of isotropic 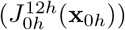 and anisotropic 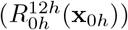 deformation (*SI Sec. 2*). Black (white) crosses indicate regions of isotropic expansion (shrinkage) with *J* > 1 (*J* < 1), where deformation is primarily isotropic. For example, tissue expansion outside Repeller 1 reduces fluxes in all directions, and a highly compressive region in front of Repeller 2 enhances diffusive fluxes in all directions. Interestingly, this region gives rise to Henson’s Node. Regions where crossed bars differ in length or color indicate 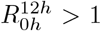 Regions where both bars vanish remain undeformed (*J* = *R* = *λ* = _1_*λ* = 1).

Comparing 4F with 4B,D reveals the cumulative effects of morphogenesis on ELCs, as predicted from the DM. Repeller 1 effectively blocks communication between blue and magenta source patches and cells inside the embryo. The yellow patch experiences reduced lateral cell-cell communication and an enhanced one along the A-P axis, overlapping with the green ELC (Fig. 4F). Green and cyan cells are both positioned near the domain of attraction, whose contraction expands their ELCs superlinearly to reach other cells brought near the PS. However, because Repeller 2 forms between them, green and cyan ELCs do not overlap (Fig. 4F). This is due to reduced diffusive fluxes across repellers, which have compartmentalizing effects. Consistent with Fig. 2J-K, the mediation of morphogen transport by attractors and repellers likely influences cell fate induction and bifurcation. It is key to notice that the modulation of ELCs by tissue deformations can be inferred from **C**—computable from experimental **v**—without solving any advection-reaction-diffusion equations.

ELCs in Figure 4G (3D view of Fig. 4F) should be contrasted with ELCs for static tissue patterning with identical patterning parameters (Fig. 4H). Without tissue flows (**v** = **0**), ELCs expand sublinearly and isotropically by simple diffusion. Instead, morphogenesis causes source cells’ signals to reach a markedly different set of cells than in static tissues. Without morphogenetic movements, blue and magenta ELCs would penetrate into the embryo, yellow and green ELCs would not communicate, instead green and cyan ELCs would overlap substantially. Crucially, morphogenesis alone does not carry any signals from cell to cell and setting *D* = 0 leads to non-expanding ELCs (Fig. 4I). We emphasize that these results are universal for any non-cell-autonomous morphogen transport mechanisms that can be modeled by an effective diffusion as in Eq. (1) (e.g., spreading via cytonemes, transcytosis). See SFig. 5 for results with other plausible patterning parameters.

### Per-cell morphogen exposure

So far, we considered *c* (# molecules/ unit area). However, when cell density (*n*, # cells/unit area) is heterogeneous, cell responses can depend on *n*[31, 32, 68, 69]. To this end, we define *q* = *c*/*n*—the average per-cell morphogen exposure (# molecules/ cell) within a tissue patch. A confluent epithelium can be compressible due to cell divisions (rate *k*_*d*_(**x**_0_, *t*)), extrusions (rate *k*_*e*_(**x**_0_, *t*)), or tissue stretching. Using continuity equations (*SI Sec. 5*), the dynamics of *n*(**x**_0_, *t*) within a Lagrangian tissue patch obeys

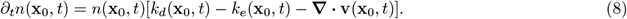

In growth-dilution models of patterning [8, 70, 71], for example, *k*_*e*_ = 0 and **∇** · **v** = *k*_*d*_ (tissue growth) so that ∂_*t*_*n* = 0 (*SI Sec. 5*). In chick gastrulation (Fig. 4), instead, uniform division, localized extrusion, and heterogeneous tissue deformation cause *n* to nearly double in the constricting embryo [65] while *n* is halved in the extraembryonic tissue [72]. It is instructive to decompose changes in *n* into changes in *i*) the patch’s number of cells *N* and *ii*) the patch’s area *A*, as *n* = *N*/*A*. Cumulative changes in 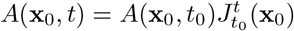 are determined by the velocity while divisions and extrusions determine cumulative changes in *N* (**x**_0_, *t*) along trajectories:

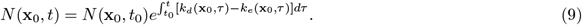

Combining Eqs. (8-9) with Eq. (6) yields Eq. (10), for *q*(**x**_0_, *t*) (*SI Sec. 5*), which explicitly accounts for division and extrusion and their role in morphogen dilution and redistribution of unbound morphogens. As in Eq. (6), non-cell-autonomous changes in *q* are still determined by 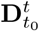 and 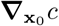, which decomposes into gradients in *q, N*, and 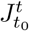. This further clarifies how morphogenesis contributes to cell-cell morphogen transport: *i*) by modulating 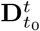 and *ii*) via heterogeneous isotropic deformation 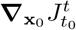.

To visualize each effect, consider an undeformed 1D tissue with uniform initial *A*(**x**_0_, *t*_0_), *q*(**x**_0_, *t*_0_), and *N* (**x**_0_, *t*_0_) and assume morphogens are conserved (no sources or sinks). Suppose that by a later time *t* tissue deformation 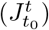 gradients develop due, for example, to heterogeneous active stresses or externally imposed forces. This deformation (morphogenesis) generates heterogeneous *q*(**x**_0_, *t*) (Fig. 5B) from uniform initial *q*. Alternatively, suppose that heterogeneous *k*_*d*_ − *k*_*e*_ develops (Eq. (9)). Heterogeneous division and extrusions likewise generate heterogeneous *q*(**x**_0_, *t*) (Fig. 5C), also from uniform initial *q*(**x**_0_, *t*_0_) (Fig. 5A). These simple examples highlight the additional generative role of morphogenesis in patterning enabled by explicitly accounting for cell density dynamics. Finally, *n*(**x**_0_, *t*) may affect patterning parameters directly. For example, both patch-level production and removal may increase with *n* (*SI Sec. 5*). Similarly, cell-cell boundaries (with density proportional to *n*) may impede transport, thereby decreasing *D*(**x**_0_, *t*) [37].

**Figure 5:**
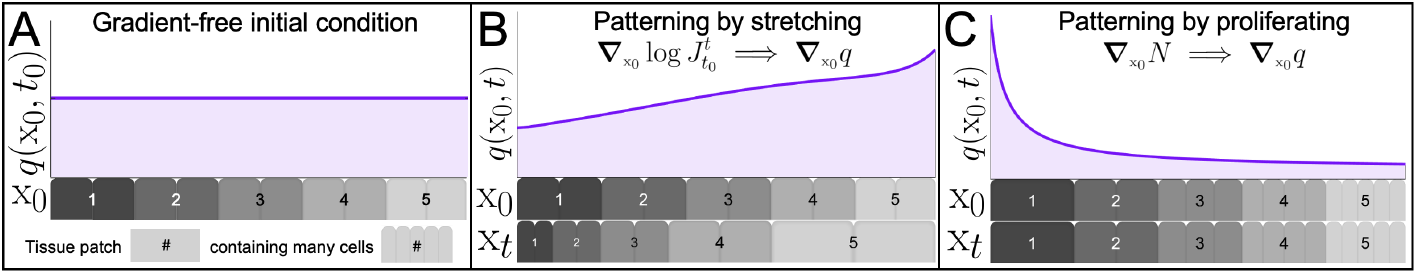
Generative patterning role of tissue deformation and cell number dynamics. A) Initially uniform *q*(x_0_, *t*_0_) and *N* (x_0_, *t*_0_) on an undeformed tissue with uniform patch areas *A*(x_0_, *t*_0_). Equivalently, *n*(x_0_, *t*_0_) = *N* (x_0_, *t*_0_)/*A*(x_0_, *t*_0_) and *c*(x_0_, *t*_0_) = *q*(x_0_, *t*_0_)*n*(x_0_, *t*_0_) are uniform. Tissue patches are differently colored and contain multiple cells (*N* (x_0_, *t*) proportional to the number of smooth polygons in each patch). B) A gradient in deformation (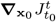, see **x**_*t*_ patch sizes) generates gradients in *q*(x_0_, *t*), while *N* (**x**_0_, *t*) remains uniform and constant as *k*_*d*_ = *k*_*e*_ = 0. Diffusion (with no-flux boundary conditions) lowers (raises) the per-cell morphogen exposure (*q*) of compressed (stretched) tissue patches. C) The same initial condition as in A and no morphogenetic movements 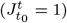, but with heterogeneous *k*_*d*_ − *k*_*e*_ generates heterogeneous *N* (**x**_0_, *t*) (contrast cells per patch with panels A and B), lowering *q* in patches with more cells.

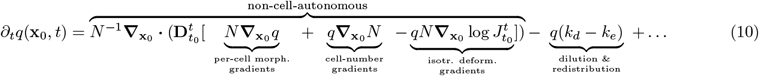

## Discussion

Morphogenesis can be thought of as a coordinate transformation—a view that dates back to D’Arcy Thompson’s seminal work [73]. This view, influential in studying how organismal form emerges and diverges, has not been adopted in the study of morphogen patterning. Yet, patterning occurs concurrently with morphogenesis in early embryos, where cells decide their fates processing morphogen exposure as they move—i.e., in their moving frames (Fig. 1).

By recasting patterning equations (advection-reaction-diffusion) in the cell frame Eqs. (2-3), we uncovered elegant connections between morphogen patterning and morphogenetic movements: tissue deformation modulates intercellular (non-cell-autonomous) diffusive morphogen transport in remodeling tissues (Fig. 2). This modulation is highest on the Dynamic Morphoskeleton [38]: a set of robust multicellular attractors and repellers that reduce complex, noisy cell trajectories to their Lagrangian kinematic units [38, 42, 44, 45]. Repellers (attractors) act as barriers (enhancers) to diffusive transport, affecting the cell-cell interaction ranges across them (Figs. 2, 4). These findings clarify how morphogenesis mediates morphogen exposure and cell-cell transport in dynamic tissues, with repellers aiding compartmentalization and fate bifurcations (Figs. 2J, 4)—consistent with recent experiments in zebrafish [42] and chick embryos [44]—and attractors supporting cell fate induction (Figs. 2K, 4). Generating sharp repellers in dynamic tissues ensures sensitivity (sharp 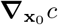, Fig. 2J) and robustness, as attractors and repellers are the most robust features in spatiotemporal flows [38], resolving a common static-tissue trade-off [74].

To assess when morphogenesis affects patterning, we defined two nondimensional numbers (Eq. (7)), computable from the experimental tissue velocities and known or candidate signaling mechanisms. These numbers account for cumulative tissue deformations in remodeling tissues and are typically distinct from the classic Péclet number. Additionally, Lagrangian coordinates (Fig 1) have another key advantage: they disentangle the confounding effect of motion in patterning [5–7], providing an equivalent static (no advection) patterning problem that encodes cells’ deforming environments into an effective diffusion tensor Eqs. (2-3). Morphogenetic movements can dramatically reshape cell-cell interaction ranges compared to static tissue patterning and Embryological Light Cones formalize these dynamic interaction ranges within highly remodeling tissues (Figs. 3, 4). In the avian gastrulation example (Fig. 4), the strong cell-cell interaction range enhancement perpendicular to the PS might aid in continuous mesoderm induction [51, 66, 75]. Repeller 1’s compartmentalizing effects resonate with the extraembryonic region’s waning regulatory ability [76–78]. Repeller 2’s compartmentalizing effects resonate with the diverging gene expression of anterior and posterior sections of the PS [38], which ultimately give rise to different cell types [79]. Finally, we incorporated cell number density dynamics into our framework (Eq. (10) and *SI Eq. 25*), formalizing key cellular mechanisms for morphogen dilution, redistribution, production and removal [50]. This framework suggests additional generative roles of morphogenesis in patterning, forming stable morphogen gradients through heterogeneous tissue deformation (Fig. 5B) or proliferation (Fig. 5C) from an unpatterned initial condition.

Historically, embryologists have compared fate maps 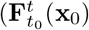, where cells normally go and what they become) with specification maps (what cells **x**_0_ would become if isolated at *t*_0_) to classify development as more ‘mosaic’ (autonomous fates decisions) or ‘regulative’ (cell-cell interactions required) [80, 81]. Our framework helps rationalize regulative development in dynamic tissues, revealing on **x**_0_ what mechanisms would enable cells to interact amid morphogenetic movements. As next steps, we plan to examine 3D morphogenesis examples, where attractors and repellers are two-dimensional surfaces. Given the difficulty of relative motion between cells and the ECM, morphogenesis can strongly mediate patterning in 3D, where the confounding effect of motion is higher but resolved in Lagrangian coordinates. Future extensions include *i*) considering cell motion’s random, mixing components [82, 83], which may affect patterning; *ii*) incorporating the timing of different signaling processes and variable competence, affecting which cells can respond to which signals; and *iii*) explicitly accounting for dynamic neighbor exchanges at single-cell resolution (as opposed to the continuum description adopted here).

Nonlinear dynamics offers the mathematical frameworks for describing spatiotemporal processes along trajectories. In cell-fate dynamics, cell trajectories live in high-dimensional gene expression space. Siggia, Briscoe and coworkers [84–86], using advanced nonlinear dynamics concepts [87], devised a striking, structurally stable compression (with minimal fitting parameters) of high-dimensional gene expression trajectories into low-dimensional cell fate dynamics between discrete attractors (stable cell states), saddle points and repellers (unstable cell states). There, morphogen exposure serves as a control parameter, affecting the routes between discrete cell states via bifurcations (reviewed in [88]). Complementary to this perspective, the Dynamic Morphoskeleton [38] compresses tissue-space cell trajectories into minimal discrete units: positional attractors (where cells converge within a domain of attraction) and repellers (where cells separate). Both techniques provide geometrical insights that do not require detailed simulations or depend on molecular details. While these dynamical systems are in different spaces (gene expression and position), they are intimately related because cell positions affect morphogen exposure, which instructs cell differentiation. This work is a step towards uniting these spaces. Connecting attractors and repellers in position space and their dual objects in the fate decision space via morphogen exposures along cell trajectories (present work) requires new ideas.

## Acknowledgements

We acknowledge Kees Weijer and Guillermo Serrano Najera for providing the experimental data used in Fig. 4 and for helpful discussions. We also acknowledge Boris Shraiman and Sreejith Santhosh for their insightful comments and Fridtjof Brauns for suggesting SFig. 3. MS acknowledges support from NSF PHY-2413073. AP acknowledges support from NIH training grant T32-GM127235.

## Author Contributions

M.S. designed research; A.P. and M.S. performed research and wrote the paper.

## Supplementary Information

### S1 Derivation of dynamic tissue patterning equations in Lagrangian co-ordinates

Advection-diffusion equations have been transformed from Eulerian to Lagrangian Coordinates [1, 2] to study the diffusive transport of scalars for incompressible [3] and compressible flows [4]. Here, we provide a compact derivation that will later enable us to add additional mechanisms relevant to biological patterning. Consider a three-dimensional tissue parametrized in Eulerian coordinates by **x** = [x, y, z], and denote by 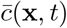 a scalar field representing the # molecules/unit volume. An overbar denotes functions of Eulerian coordinates **x** and *t*. We assume that 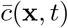 is advected (i.e., co-moves) with the tissue with velocity 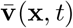. Denoting by Ω_*t*_ an arbitrary tissue region (volume) at time *t*, the rate of change of the total amount of 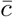 in Ω_*t*_ depends on sources and sinks of 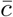 in Ω_*t*_ and the diffusive fluxes through its boundary surface (∂Ω_*t*_):

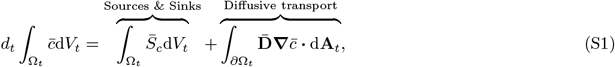

where *d*_*t*_ denotes the total (or Lagrangian) time derivative; d*V*_*t*_ an infinitesimal volume element in Ω_*t*_; d**A**_*t*_ a vector representing an infinitesimal area element of ∂Ω_*t*_ (magnitude represents its infinitesimal area with a direction outward perpendicular to the area element); 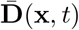 the diffusion tensor; and 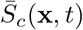 sources and sinks of 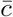. We keep a general 3D notation and note that for 2D tissues, Ω_*t*_ represents an area and d**A**_*t*_ its boundary curve. For 2D tissues (i.e., confluent monolayer), 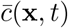 can represent # molecules/unit area (Fig. S1). When the advective velocity of 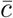 differs from that of the tissue, (S1) can be modified to account for advective fluxes due to their relative velocity (see following subsections).

**Fig. S1:**
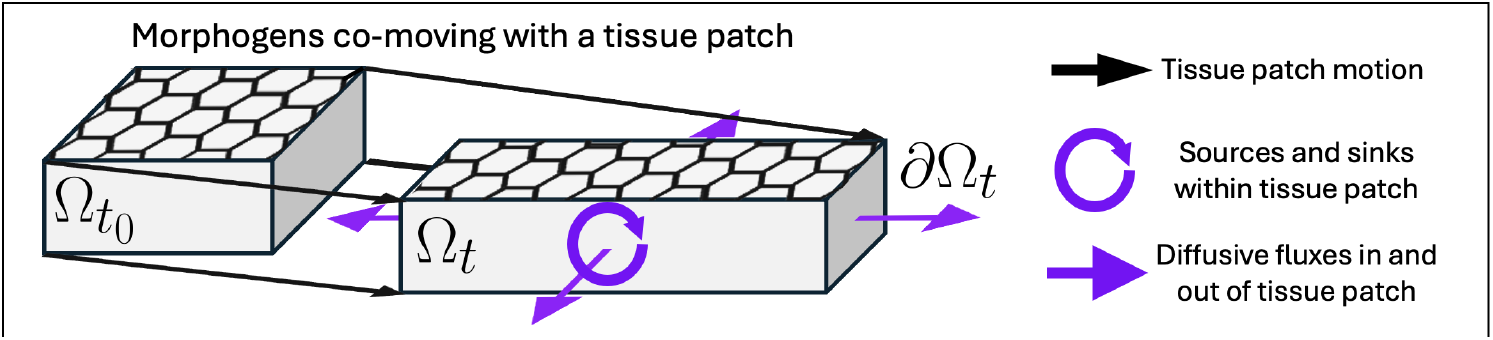
Morphogen exposure in a moving tissue patch. Initial tissue patch (material volume 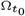 containing cells and ECM) and the same tissue patch at a later time (material volume Ω_*t*_), after moving and deforming. Arrows represent the two terms on the right-hand side of (S1), including the production and removal of morphogen within the tissue patch and diffusive transport to and from neighboring patches through lateral patch boundaries.

#### S1.1 Transformation into Lagrangian coordinates

We transform (S1) from Eularian Coordinates **x** into Lagrangian Coordinates **x**_0_ (the cell frame) using the trajectory map 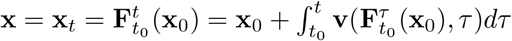. The left-hand side of (S1) transforms as

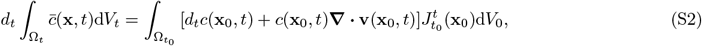

where 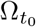 denotes the tissue volume at the initial time *t*_0_ with infinitesimal volume elements d*V*_0_ and 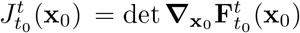, with 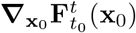 denoting the deformation gradient, i.e., the Jacobian of the trajectory map with respect to **x**_0_. In (S2), functions without 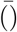 represent the same Eulerian function evaluated along tissue trajectories (for example, 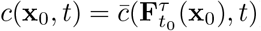). To obtain (S2), we used 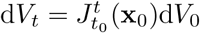 and

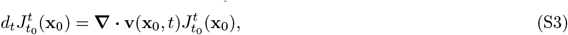

which can be derived by Liouville’s theorem [5] and relates the Lagrangian (i.e. cumulative along trajectories) rate of change of a tissue volume initially centered at (**x**_0_, *t*_0_) with the instantaneous velocity divergence at the current position 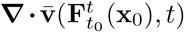, which in Lagrangian coordinates is denoted by **∇**· **v**(**x**_0_, *t*). The first term on the right-hand side of (S1) transforms as:

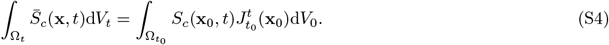

The second term, representing diffusive flux through Ω_*t*_, requires us to transform both the gradient and d**A**_*t*_. The former, 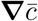, that is 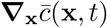, becomes 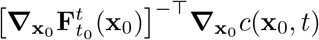 (^⊤^ denotes matrix transpose) and can be obtained from the chain rule using the trajectory map. The latter transforms as 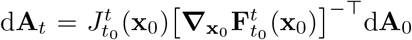 [5], and relates the boundary area element d**A**_*t*_ of ∂Ω_*t*_ to the same area patch d**A**_0_ of 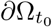. Rearranging terms across the inner product yields

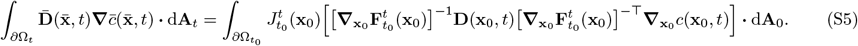

Putting together Eqs. (S2, S4, S5), transformed back into a volume integral by the divergence theorem, we have:

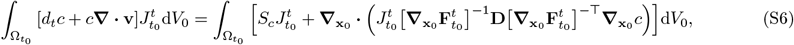

where all functions are in Lagrangian coordinates (i.e., with no 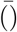), and hence we omit their dependence on **x**_0_ and *t* whenever possible. Converting (S6) into differential form yields

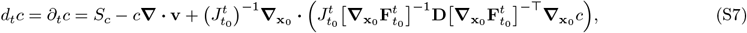

where *d*_*t*_*c*(**x**_0_, *t*) = ∂_*t*_*c*(**x**_0_, *t*) as there is no advective derivative in the frame moving with cells. If we further assume isotropic Eulerian diffusivity **D**(**x**_0_, *t*) = *D*(**x**_0_, *t*)**I** in (S1), (S7) simplifies to

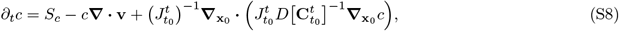

recovering Eqs. (2,3,6) in the main text. From the Cauchy-Green strain tensor definition 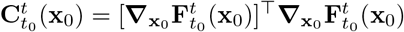, it follows that 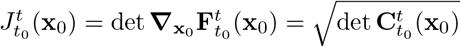

#### S1.2 Instantaneous limit

To gain intuition, we take the limit *t* →*t*_0_ to examine the first-order correction (in time) to diffusive fluxes in the cell frame. The right Cauchy-Green strain tensor becomes [6]

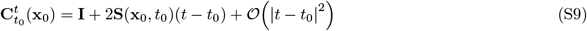

in terms of the Eulerian rate of strain tensor 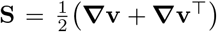 Substituting (S9) into (S8) yields 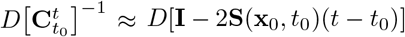. Similarly, 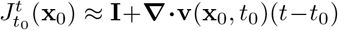. Hence, keeping leading order terms in *T* = |*t*−*t*_0_|, (S8) yields:

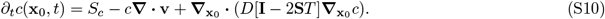

showing that the leading order effects on morphogen transport in the cell frame are governed by **S**(**x**_0_, *t*_0_) and are maximal along its orthogonal eigenvectors with modulation encoded in its eigenvalues *s*_2_(**x**_0_, *t*_0_) ≥ *s*_1_(**x**_0_, *t*_0_). Over short times, the correction to *D***I** becomes additive instead of multiplicative and is typically negligible in multicellular flows because *T* ≈ 0 and strain rates (**S**) are small.

#### S1.3 Relative tissue-ECM velocities

Here we briefly summarize extensions of our approach to account for relative motion between cells and the ECM. Suppose that cells and ECM move with distinct velocity fields 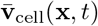 (for cells) and 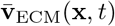. Note that the Lagrangian coordinates of interest **x**_0,Cell_ (the cell frame) are now distinct from **x**_0,ECM_. We will use the shorthand **v** = **v**_cell_ and **x**_0_ = **x**_0,Cell_. We need *c*(**x**_0_, *t*) to address how *i*) motion affects morphogen exposure histories and *ii*) deformation affects cell-cell interaction ranges. Returning to (S1), an additional advective flux term 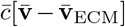 arises due to the mismatch between the motion of the ECM (containing morphogens) and the cells (Ω_*t*_). Carrying this through the derivation yields the following additional term on the right-hand side of the differential equations (S8):

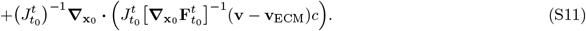

Note that **v** and **v**_ECM_ here are both taken as functions of **x**_0_, representing cells’ own velocities along trajectories and the velocities of ECM at the corresponding spatial coordinates. Note also that 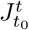 and 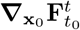 encode deformation associated with the cell velocities, as we have transformed into their Lagrangian coordinates, not those of the ECM. Accounting for the correction term ((S11)), consider the following cases:

1. If **v** ≠ **v**_ECM_, cells experience an effective advective flux due to their motion relative to the morphogen concentration. In general, we have

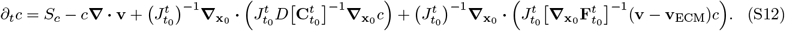
2. If **v** = **v**_ECM_, we recover (S8).
3. If cells move relative to a fixed ECM (e.g. ECM acting as a fixed substrate), (S12) reduces to

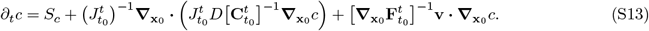
4. If ECM flows relative to a static tissue, **x**_0_ = **x** and 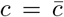 (S12) reduces to an ordinary advection-diffusion equation:

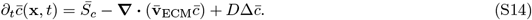

### S2 Characterization of Lagrangian deformations

In (S8), the diffusive flux depends on the tensor **C** (dependence on *t*_0_ < *t* and **x**_0_ as above). **C** can be diagonalized in its eigenbasis as **C** = **VΛV**^⊤^ where **Λ** is a diagonal matrix containing the eigenvalues *λ* ≥ _1_*λ* > 0 and **V** is a matrix whose columns are the corresponding eigenvectors (***ξ*** ⊥ _1_***ξ***). In one dimension, we trivially have **C** = *λ*. In two dimensions, eigenvalues and eigenvectors describe the local direction and magnitude of the highest and lowest Lagrangian strain (Fig. S2A). Alternatively, one can decompose deformation into isotropic and anisotropic components (Fig. S2B)

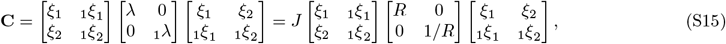

where 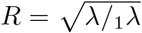 represents the deformation anisotropy (i.e., the deformed ellipse aspect ratio) and 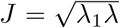 the isotropic deformation (i.e. the ratio of the deformed and undeformed patch area). Note that in 3D, one can still extract 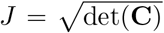 (volumetric strain), but that there are instead multiple degrees of freedom for characterizing the anisotropy of the deformation. For any time interval [*t*_0_, *t*], there may exist particular tissue patches **x**_0_ where 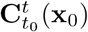 does not have distinct eigenvalues (in 2D: 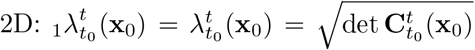). These distinct locations mark regions of purely isotropic deformations 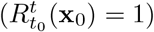, sometimes referred to as 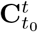 singularities [7] (Fig. S2B).

**Fig. S2:**
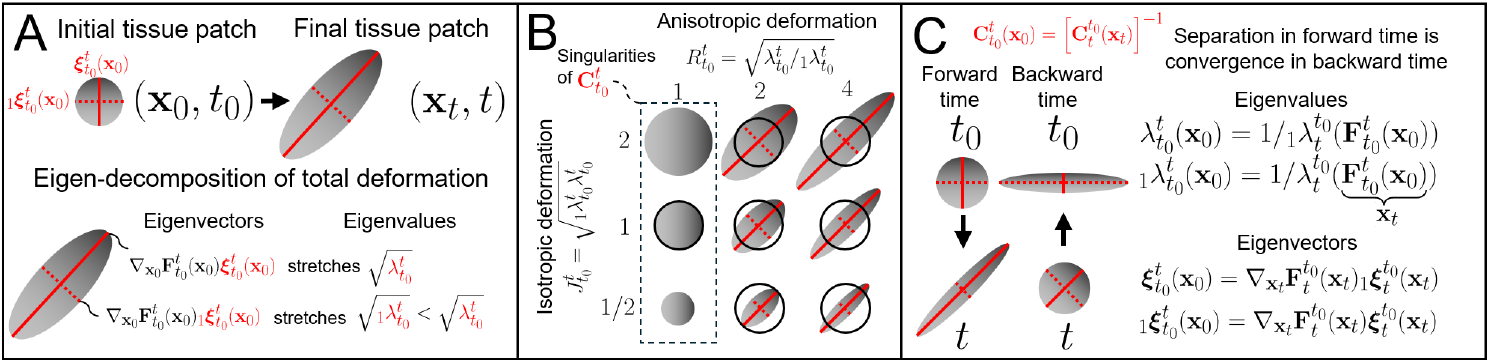
Lagrangian deformation of a tissue patch. A) Lagrangian 2D tissue patch deformation quantified by 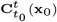. B) Alternative decomposition into isotropic and anisotropic deformations. C) Attractors and repellers are computed using forward and backward time trajectories (see e.g., [8]). Deformations of a tissue patch can be displayed at initial and final embryo configurations. They can be computed from forward and backward time analysis, are directly related and computed from a single computation [9].

Finally, it is instructive to know whether a particular tissue patch **x**_0_ experienced only shrinking or stretching deformations. This information is readily available from the _1_*λ* (least stretching in the direction _1_***ξ***) and *λ* (greatest stretching in the direction ***ξ***). Regions where _1_*λ* > 1 indicate that all directions stretched. Similarly, regions where *λ* < 1 indicate that all directions shrank. In incompressible flows, these regions cannot exist as _1_*λ* = 1/*λ* (*J* = 1 in Fig. S2B).

### S3 Quantitative criteria for dynamic tissue patterning

In practice, a developmental process starts at an initial time *t*_0_ and data is collected over a duration *T* of interest for patterning. For example, in the problem of mesoderm induction during chick gastrulation (main text Fig.4), one can set *t*_0_ = *HH*1 (a developmental stage) and *T* = 12*h* (the typical duration of gastrulation).

#### S3.1 Nondimensional parameter Ω_1_

Over [*t*_0_, *t*_0_ + *T*], the largest finite-time Lyapunov exponent (in short, FTLE) is defined as

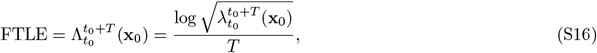

where 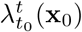 is the largest eigenvalue of 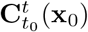. To understand its physical meaning, start with the linear, time-independent convergent extension flow 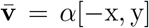, *α* > 0 (main text Eq. (4)). In time-independent flows, it is standard to set *t*_0_ = 0 as trajectories, **C**, FTLE etc., depend only on the time interval length *T* and not on the specific initial time *t*_0_. For this flow 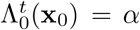, meaning that any circular patch of diameter *d*_0_ will maximally stretch along y into an ellipse with major diameter *e*^*αT*^ *d*_0_. However, there is no simple connection between the FTLE and the velocity strain rates for nonlinear and time-dependent flows. To gain intuition in these flows, without loss of generality, fix *t*_0_ = 0 and focus on a single tissue patch labeled by its initial condition **x**_0_. 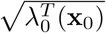 quantifies the maximum stretching of two points near **x**_0_ over the time interval [0, *T*] (Fig. S2A). However, this cumulative stretching is typically not simply a *constant* \**T* but a complicated *f* (*T*) (and initial time *t*_0_). Yet, it is instructive to quantify the time-averaged stretching rate in the direction of maximum stretching for a tissue patch starting at **x**_0_. The FTLE quantifies precisely this rate by implicitly supposing that the cumulative stretching 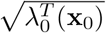 was achieved by an equivalent linear flow: 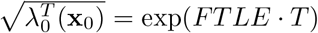. Taking the logarithm of this relation gives (S16). Therefore, for any **x**_0_ and time interval 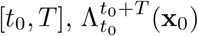 quantifies the (time) average maximum stretching rate of a tissue patch starting at **x**_0_, overcoming the complications caused by nonlinearities and temporal dependence of 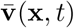 to provide similar insights available for linear, time-independent velocities. To probe temporal heterogeneity, one can always explore different *t*_0_ and *T*.

A related, useful question is when (i.e., at which *t* ∈ [0, *T*]) and where (**x**_0_) will deformations become appreciable? To gain intuition, we start again with the linear example above, where the maximum separation of the initially *d*_0_-close nearby points obeys 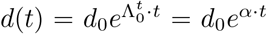. For 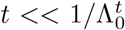 (i.e. *t* << 1/*α*), *d*(*t*) ≈ *d*_*0*_. By contrast, for *t* >> 1/*α* we have *d*(*t*) >> *d*_0_, meaning that there is a characteristic time 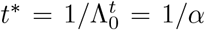 after which separation increases appreciably. For nonlinear, time-dependent velocities, this characteristic time is called

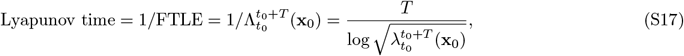

and is computable for any spatiotemporal flows from experimental tissue velocities [8] or sparse, noisy cell trajectories [10]. Intuitively, the Lyapunov time can be thought of as a *deformation time scale*, revealing where and when deformations in the tissue become appreciable. Putting together Eqs. (S16-S17), we define the nondimensional parameter

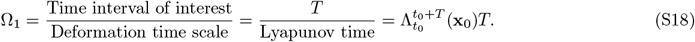

In practice, one can set i) *t*_0_ = 0 as the beginning of the experiment or the initial developmental stage for the problem under study and ii) *t* = *T*, where *T* is the time interval of interest for patterning. For example, in the problem of mesoderm induction during chick gastrulation (main text Fig. 4), one can set *t*_0_ = 0 = *HH*1 stage and *T* = 12*h*. Then, one computes 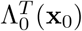 from experimental velocities [8], from which (S18) reveals a spatial map (Fig. S4C) of the tissue patches **x**_0_ where deformations are appreciable (Ω_1_ > 1, i.e. time scale of interest > deformation time scale).

For compressible flows, *λ* and _1_*λ* can vary independently, meaning that the tissue could converge more in one direction than it extends in the other direction, or even converge in every direction, so that the strongest deformation is associated with convergence, not separation. In these cases, one should consider in (S17) and (S18) both the forward time FTLE ((S16))—capturing maximum stretching and defining repellers—and the backward time FTLE for the same patch: 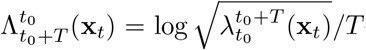 —capturing maximum convergence and defining attractors—and use whichever FTLE is larger, i.e., resulting in the smallest Lyapunov time. This ensures that the Lyapunov time (and therefore Ω_1_) is always positive and accounts for the highest stretching or compressive deformation. Note that the backward time FTLE can also be directly computed from the smallest eigenvalue 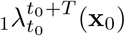 of the forward time 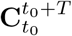 (Fig. S2C) and represents compression along 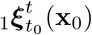 (see Algorithm 1 for the precise formula).

Ensuring that deformations become appreciable during [*t*_0_, *t*_0_ + *T*] guarantees that 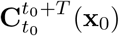 deviates from **I**, implying 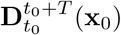 deviates from *D***I** (main text Eq. (3)), i.e. the equivalent diffusion tensor in the cell frame is distinct from the standard Eulerian diffusion tensor. This constitutes condition *i*) in main text Eq. (7). See Fig. S4 for Ω_1_ in avian gastrulation flows and in the 1D SDD model used in Fig. 2J-K.

#### S3.2 Nondimensional parameter Ω_2_

Inspection of (S8) reveals that for diffusive fluxes in the cell frame to be affected by deformations, in addition to ensuring 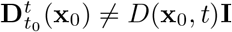 (guaranteed by condition *i*): 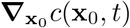 should remain nonzero until deformation effects become appreciable. Generally, 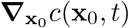 depends on boundary conditions, source, sink, and reaction terms. Hence, we distinguish two cases. If there are mechanisms to sustain 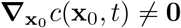, then condition *i*) is the only one needed for morphogenesis to affect patterning. This is the case, for example, in the SDD model discussed in the main text Fig. 2 I-K. By contrast, if there are no mechanisms to sustain 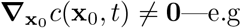—e.g. because a morphogen was locally injected into a tissue—one needs to ensure that diffusion does not flatten gradients in the patterning region of interest before deformations become appreciable, leading to the second nondimensional parameter

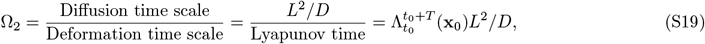

where *L* is the size of the patterning region of interest, and *D* is a characteristic diffusivity. Here too, for compressible flows one should use the larger of 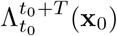 and 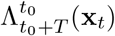 for each tissue patch (See Algorithm 1). Ω_2_ > 1 ensures that before diffusion flattens 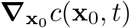, deformations effect have become appreciable and affected cell-cell diffusive fluxes. If diffusion is not the fastest mechanism to flatten 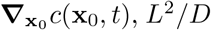 should be replaced with the corresponding time scale. For example, if degradation is faster than diffusion and there are no mechanisms to sustain gradients, it will drive *c*(**x**_0_, *t*) → 0 and hence 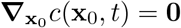.

##### S3.2.1 Relation between Ω_2_ and the Péclet number

Ω_2_ is generally different from the classic Péclet number (Pe = |**v**| *L*/*D*), a ratio of advective and diffusive transport rates. This is because, in the cell frame, advection is absent ((S7)), and its effect modulates diffusive flux due to tissue deformations 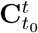. In fact, a spatially uniform velocity would have Pe = |**v** |*L*/*D* ≠ 0 but Ω_2_ = 0 because advective transport in the Eulerian frame would be nonzero, but that would not affect cell-cell transport.

However, considering the alternative Péclet number Pe = |*α*|*L*^2^/*D* based on a strain rate (*α*), Ω_2_ = Pe in two cases. First, in the case of time-independent velocities linear in space (e.g. 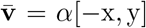 in main text Eq. (4)). In this idealized linear case, 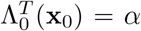 (Sec. S3.1) meaning that knowledge of the stationary and spatially constant velocity strain rate is sufficient to quantify cumulative tissue deformations over a finite time. Second, in the case of nonlinear and time-dependent **v**, only in the instantaneous limit (i.e., infinitesimally short finite time interval):

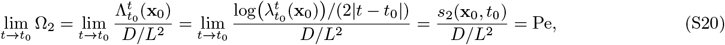

where we used that for small *t*−*t*_0_, 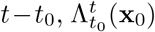 corresponds to the instantaneous, local maximum strain rate *s*_2_(**x**_0_, *t*_0_) [11], which is the largest eigenvalue of the tissue velocity rate-of-strain tensor **S**(**x**_0_, *t*_0_) = 0.5([∇**v**(**x**_0_, *t*_0_)]^⊤^ + ∇**v**(**x**_0_, *t*_0_)). Intuitively, at the initial time *t*_0_, the tissue is undeformed, and a tissue patch will initially start deforming maximally with the highest stretching rate *s*_2_(**x**_0_, *t*_0_) along the local stretching direction associated with the dominant eigenvector of **S**(**x**_0_, *t*_0_), as shown in (S10). However, the general spatiotemporal features of **v**(**x**_0_, *t*) will induce finite-time deformations of tissue patches that cannot be related to instantaneous velocity features but are precisely captured by 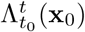 and hence Ω_1_ and Ω_2_. In summary, Ω_1_ and Ω_2_ capture the effects of finite-time tissue deformations on patterning in general nonlinear morphogenetic processes and are computable from experimental data of tissue velocities or single cell tracks [10]. These numbers reveal that patterning in dynamic tissues is intimately tied to cumulative tissue deformations instead of the classic Péclet number—relevant for static tissue patterning. For an algorithmic procedure to verify Ω_1_ > 1, Ω_2_ > 1, see Algorithm 1.

### S4 Embryological Light Cones

The idea of light cones in pattern-forming systems has been explored in simple 2D cellular automata [13, 14]. The notion of a light cone enabled quantifying the degree of self-organization (distinguishing self-organized complexity from spurious order or randomness) because it accounts for non-cell-autonomous causal influences and their time-dependent spatial extents. In these models, material points do not move and their light cones are defined by a constant, uniform propagation speed (e.g., *c* in the cosmological case). In the embryological case, information propagation rates can be non-uniform, non-constant, and anisotropic.

For a cell labeled 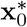, we define its past cone 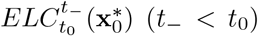 and future cone 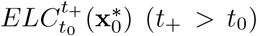 as regions that can affect the cell’s state, and regions that the cell can affect, over a specified time interval and through some specified signal propagation mechanism. ELCs must be examined in Lagrangian coordinates (i.e., **x**_0_ or **x**_*t*_) as they label sets of cells, facilitating comparisons with fate maps and specification maps. Affecting a cell can be formalized in various ways and depends on what type of information the cell responds to, e.g., integrated exposure, rate of change of exposure, or a simple threshold. Additionally, this threshold might be heterogeneous or dynamic, taking a cell’s competency to respond to signals into account [15]. For simplicity, we consider a low exposure threshold *c*_min_—a necessary condition for any signal-sensing. Choices for a signaling threshold will affect ELC extents and are both necessary—because a diffusing *c*(**x**_0_, *t*) is a continuous field that becomes nonzero everywhere after a finite time—and biologically important as signaling quantities (e.g. molecules) are discrete and perceptual thresholds are finite.

#### Algorithm 1 Procedure to verify if morphogenesis mediates morphogen transport (main text, Eq. (7))

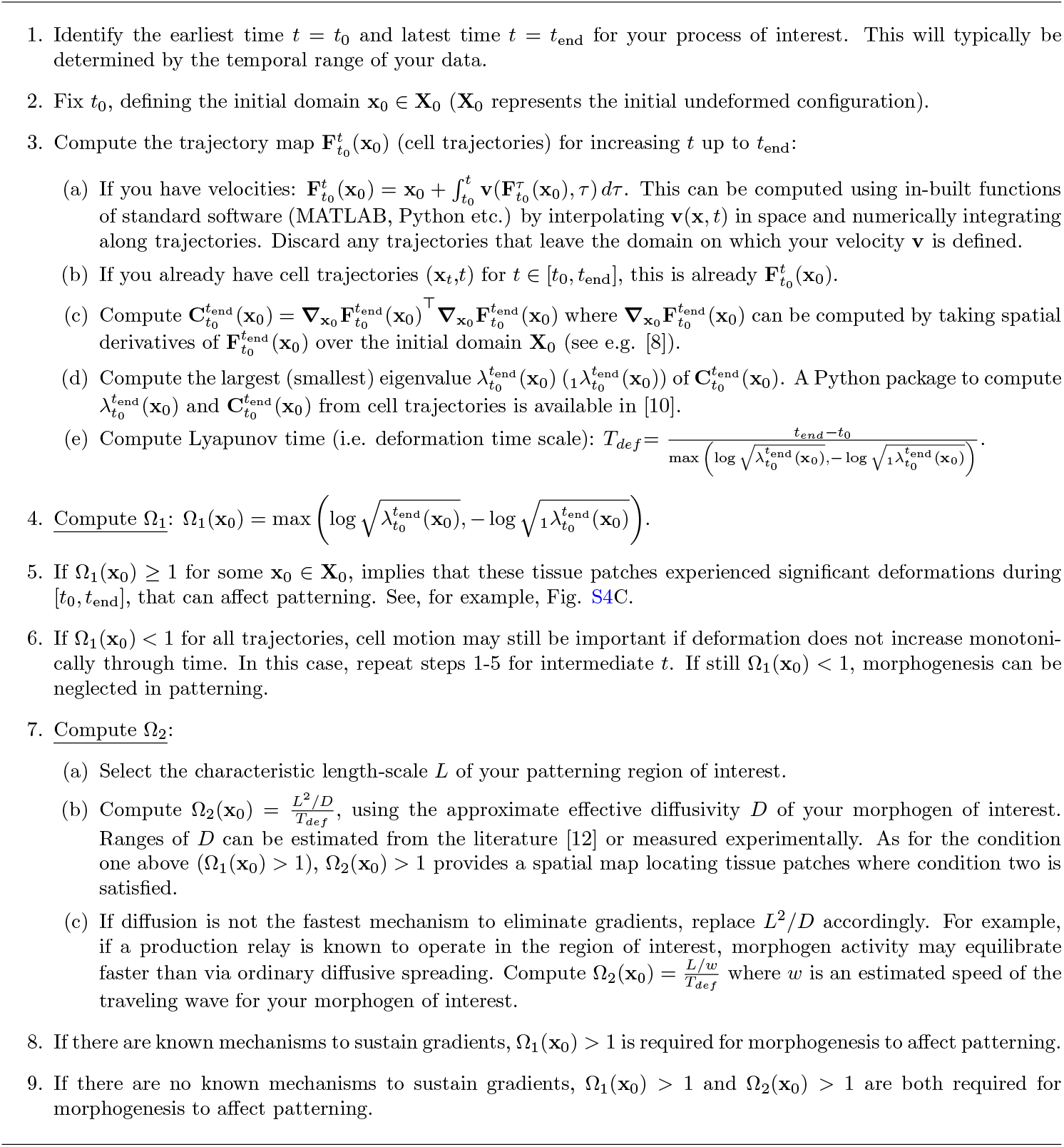

For simple static tissue patterning (**x** = **x**_0_), some ELCs can be derived analytically in 1D or 2D. Consider a fixed diffusivity *D* and the following common scenarios where a cell at 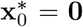 emits a signal onto a semi-infinite domain for all *t* ≥ *t*_0_ = 0:

1. **Synthesis and diffusion (1D)**: ∂_*t*_*c* = *D*Δ*c*, with flux *Q* from the boundary has analytical solution 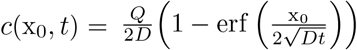, and therefore 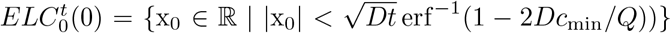, expanding sub-linearly. For visualization of simulated 2D examples, see main text Fig. 3A left and Fig. S5A.
2. **Synthesis, diffusion, and degradation (1D)**:(∂_*t*_*c* = *D*)Δ*c* − *kc*, with flux *Q* from the boundary has analytical steady state 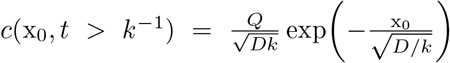, and therefore 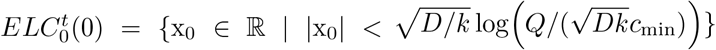 for all *t* > *k*^−1^, the approximate time for steady state formation when *k* > *D*/*L*^2^ (i.e. with significant degradation), expanding up to a fixed radius. For visualization of simulated 2D examples, see main text Fig. 3A right and Fig. S5C.
3. **Production relay (1D)**: ∂_*t*_*c* = *D*Δ*c* + *βc*(*c*_max_ − *c*), with positive feedback parameter *β*, results in a traveling wave with speed 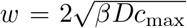 [16] away from the initial source at 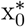. If there is an Allee effect, so that a finite amount of morphogen is needed to trigger positive feedback, we have ∂_*t*_*c* = *D*Δ *c* + *βc*(*c*_max_ − *c*)(*c* − *a*) with activation threshold 0 < *a* < *c*_max_/2, and the wave speed becomes 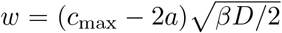 [17]. In both cases, once the front has stabilized (*t* >> (*βc*_*max*_)^−1^) we can approximate 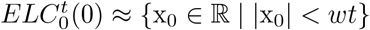, expanding linearly. For visualization of simulated 2D examples, see main text Fig. 3A middle and Fig. S5B.

Estimating a past cone 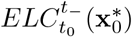 requires determining the set of points **x**_−_ for which a given 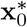 lies in their future light cone 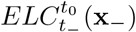. Unlike rewinding tissue patch trajectories 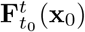 (a reversible mapping), one cannot simply run a diffusive process in reverse. For static patterning schemes, the past cone will always be the mirror image of the future cone because communication solely depends on the time duration (*T*) and the (fixed) distance between the sender and receiver (PDE for *c*(**x**_0_, *t*) is autonomous). In dynamic tissues, time-symmetry is broken by the explicit time-dependence of cell-cell separation (PDE for *c*(**x**_0_, *t*) becomes non-autonomous). For these cases, simulations will generally be required to determine ELCs. But, as we have emphasized, whether tissue deformation causes ELC expansion to accelerate, decelerate, or become anisotropically directed can be qualitatively understood from 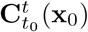 and the Dynamic Morphoskeleton (see main text Fig. 4), without solving advection-reaction-diffusion equations or measuring patterning parameters (e.g. *D, k, β*) experimentally. Finally, ELCs can be adapted to any information propagation mechanisms. For example, if cell fates respond to mechanical signals via mechanotransduction, one might consider mechanical information propagated via elastic stress-strain relationships and estimate their speed limits [18] from tissue mechanics.

### S5 Per-cell morphogen exposure in the Lagrangian frame

To derive per-cell morphogen exposure (# morphogens/cell) in the Lagrangian frame, we account for cell density dynamics, i.e., the number of cells packed into each tissue patch over time. We consider a 2D cell monolayer as a continuum composed of cells as mass carriers. From standard mass transport, a continuum with density 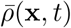 (mass per unit area) obeys the continuity equation

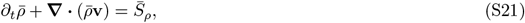

where 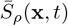 models sources or sinks of mass (mass rate per unit area). To connect this to the cell number density (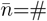 cells per unit area), we assume that cells have, on average, a fixed mass *m*_cell_, or, equivalently, a fixed average volume and physiological density. Then, we have that 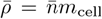, which substituted into (S21) yields a standard continuity equation for cell number density [19, 20]:

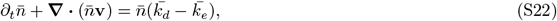

where 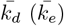 represents division (extrusion) rates per cell. Following the approach in Sec. S1, (S22) can be written in Lagrangian coordinates as

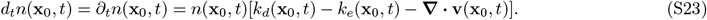

Next, we define the number of morphogens per cell *q* = *c*/*n*, i.e. the allocation of morphogen exposure to cells occupying the same tissue patch (*c* and *n* both defined per unit area). Using (S23) and substituting *c* = *nq* into (S8) gives

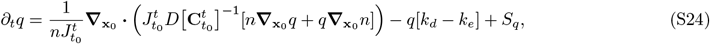

where *S*_*q*_ = *S*_*c*_/*n* is the per-cell rate of morphogen production and removal. For example, if each cell produces morphogens at rate *k*_*p*_(**x**_0_, *t*) and irreversibly removes morphogens at rate *k*_*r*_(**x**_0_, *t*) (reviewed in [21]), then *S*_*q*_(**x**_0_, *t*) = *k*_*p*_(**x**_0_, *t*) − *k*_*r*_(**x**_0_, *t*)*q*(**x**_0_, *t*). It is instructive to rewrite (S24) to represent changes in *n* due to changes to *i*) patch area *A* and *ii*) cell number *N*. By substituting 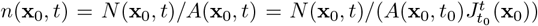 into (S24), we obtain

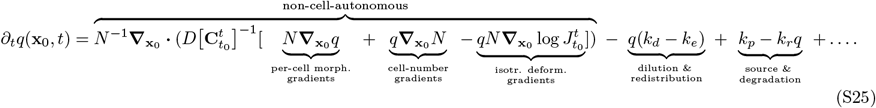

The cell number 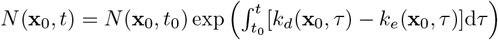 tracks the total number of cells along a patch’s trajectory, without considering how the patch area changes, which is instead tracked by 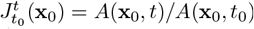, where *A*(**x**_0_, *t*_0_) is uniform, describing the initial area of undeformed tissue patches. (S25) elucidates separate effects of cell number dynamics and tissue deformations on the per-cell allocation of morphogen exposure.

#### S5.1 Simplifications with incompressibility or mass conservation

One can simplify (S25) if tissue flows are incompressible or if the total number of cells (i.e., mass) is conserved. Here we neglect production (*k*_*p*_) and removal (*k*_*r*_) terms for brevity, as they are unaffected. If the tissue is incompressible 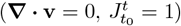, then *N* and *n* are interchangeable and (S25) reduces to

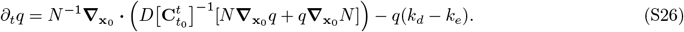

If, instead, cells numbers are conserved, *N* (**x**_0_, *t*) = *N* (**x**_0_, *t*_0_) is constant in time. If *N* (**x**_0_, *t*_0_) is also uniform in space (i.e., initial undeformed tissue patches have uniform number of cells), (S25) reduces to

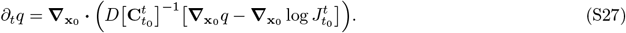

Finally, if we have both cell number conservation and incompressibility—*n*(**x**_0_, *t*) = *n*(**x**_0_, *t*_0_), *N* (**x**_0_, *t*) = *N* (**x**_0_, *t*_0_), *k*_*d*_ = *k*_*e*_–then *q* ∝ *c* and (S25) becomes

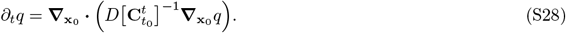

Note that (S24) and (S25) also extend to 3D tissues where 3D tissue volume elements replace 2D tissue patches. In 3D, *q* carries the same meaning, but *c* and *n* are defined per-unit-volume, *N* is the number of cells in volume element moving with cells, and 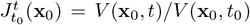 is a volume ratio. In 3D, confluent tissues, cells may die and divide, but ∂_*t*_*ρ* = ∂_*t*_*n* = 0 as cells are effectively incompressible. In this case, (S25) becomes

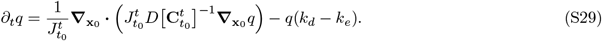

and *q* and *c* become interchangeable. (S29) recovers, as a simple example, the dynamics of existing 1D growth-dilution models [22, 23, 24, 25], which include a morphogen dilution term −*c***∇** · **v** associated with tissue growth (i.e. **∇** · **v** = *k*_*d*_, *k*_*e*_ = 0, conserving *n*). (S25) instead applies to general 2D flows, where proliferation can occur under confinement or tissue stretching, and with any anisotropic tissue deformation.

#### S5.2 ELCs for dynamic tissue patterning

When cell number density dynamics are known, describing ELCs with *q* = *c*/*n* instead of *c* better represents the signal a cell can sense. Consider the following simple dynamic tissue patterning scenarios on a semi-infinite 1D domain, assuming *k*_*d*_ = *k*_*e*_, uniform initial cell density, and uniform deformation (i.e. (S28)), and 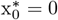, *t*_0_ = 0:

1. **Shrinking Tissue (1D)**: v = −*α*x, *α* > 0, ∂_*t*_*q*(x_0_, *t*) = *De*^2*αt*^Δ*q*, with fixed *q*_0_ at the 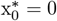 boundary has analytical solution 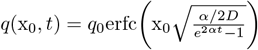, and therefore 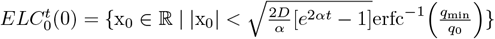, expanding at an increasing rate that becomes superlinear.
2. **Expanding Tissue (1D)**: v = *α*x, *α* > 0, ∂_*t*_*q*(x_0_, *t*) = *De*^−2*αt*^Δ*q*, with fixed *q*_0_ at the 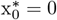 boundary has analytical solution 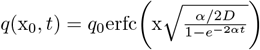, and therefore 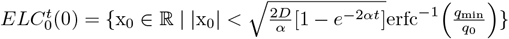, expanding at a decreasing rate and approaching a fixed radius beyond which growth outpaces diffusion.

This 1D analysis is consistent with the simulated expansion of ELCs in 2D shrinking and expanding tissues (cf. Fig. 3).

### Additional Supplementary Figures

**Fig. S3:**
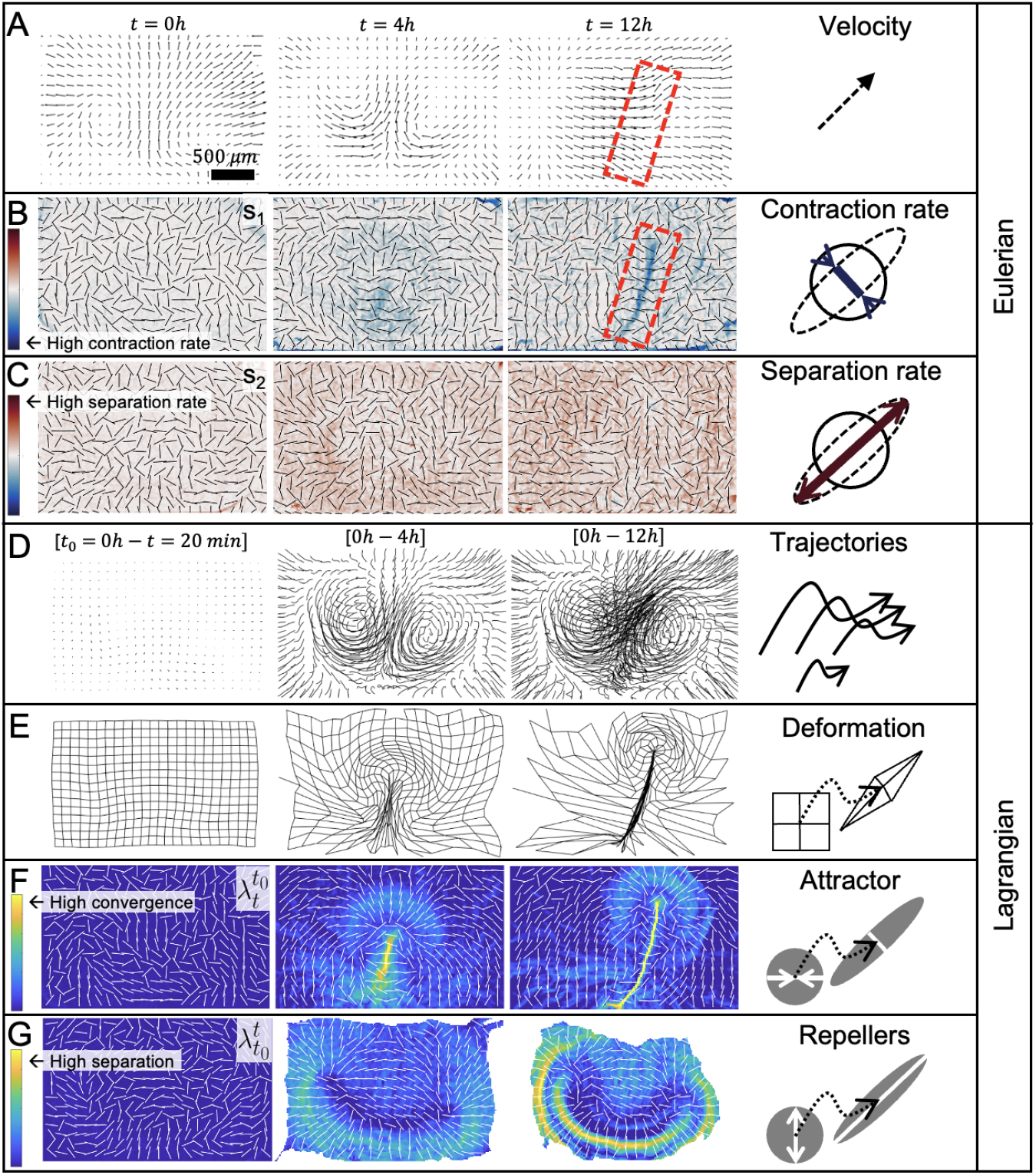
Connections between Lagrangian and Eulerian quantities. A) Chick Gastrulation velocities **v**(**x**, *t*) from *t*_0_ = 0*h* (HH1) to 12*h* (HH3) used in Fig. 4. Velocity plots fail to locate regions of convergence and separation in time-dependent flows. For example, at the Primitive Streak (red rectangle), the tissue exhibits high convergence without an obvious signature in **v**. Instead, the frame invariant rate-of-strain tensor should be considered to locate local (in space and time) regions of convergence or separation [6, 11]. B) *s*_1_ (**x**, *t*) field and directions (white bars) show the smallest eigenvalue (eigenvector) of the rate of strain tensor **S**(**x**, *t*) = 1/2(∇**v**(**x**, *t*) + [∇**v**(**x**, *t*)]^⊤^), showing the orientation and intensity of maximal local contraction rates, marking a short-time attractor at the Primitive Streak [26]. C) Same as B for local separation. Maximal separation rates are quantified by the largest eigenvalue *s*_2_ (**x**, *t*) of **S**(**x**, *t*) along its eigenvector (black bars). B shows no sign of repellers in instantaneous fields. The (Eulerian) **S**(**x**, *t*) is frame invariant yet agnostic to cell paths. Tissue patches, instead, move and integrate deformations along their trajectories (D), visualized by a deforming Lagrangian grid (E). The Dynamic Morphoskeleton (DM), composed of attractors (F) and repellers (G), precisely accounts for this cumulative deformation over a finite time. F) Largest contraction ratio 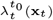 and orientation (white bar) along cell trajectories. G) Largest separation ratio 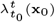 and expanding direction (white bar) along cell trajectories. Comparing E-G with B-C shows robust morphogenetic features that develop over time (Lagrangian) and are not contained in Eulerian fields.

**Fig. S4:**
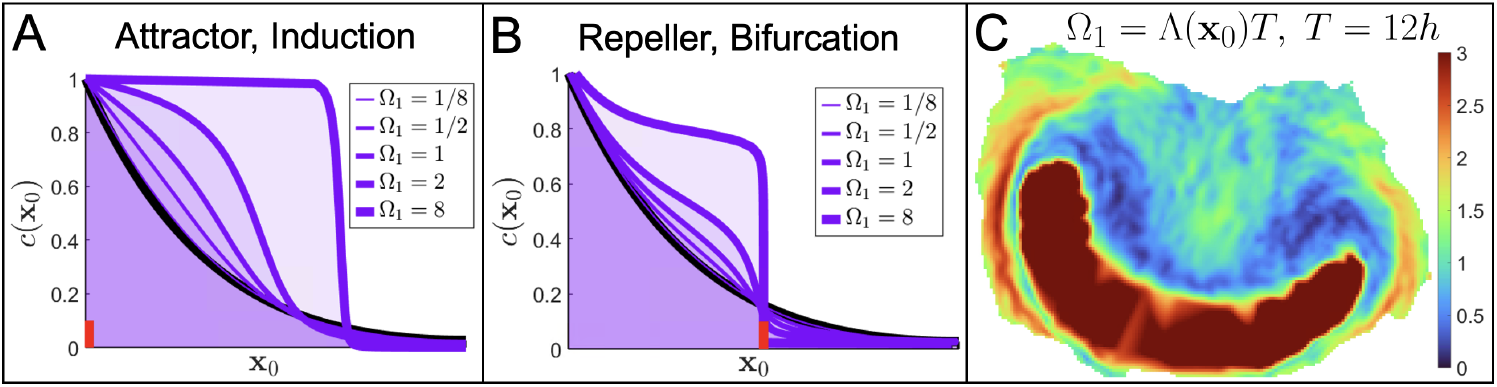
Non-dimensional Parameter Ω_1_. A-B) Same as main text Fig. 3J-K for different values of Ω_1_. Black curve marks SDD static tissue patterning (**v** = **0**), while purple curves mark the concentration profiles *c*(**x**_0_, *t*) for dynamic tissue patterning (**v** ≠ **0**) with increasing deformation rates. For Ω_1_ > 1, deformations affect morphogen exposure dynamics. A) Velocity field v(x) = *α*(x − *L*/2) exp[−(x − *L*/2)^2^/(2(0.15)^2^)]. B) Velocity field v(x) = −*α*x exp[−x^2^/(2(0.25)^2^)]. Both A and B have the largest Ω_1_ = *αT* ((S18)) on the Dynamic Morphoskeleton (attractor (A) and repeller (B) marked in red). All simulations use diffusivity *D* = 0.001, degradation rate *k* = 0.01, time *T* = 100, and domain size *L* = 1 with fixed *c*(0) on the left boundary (arbitrary units) and no flux on the right boundary. C) Ω_1_ (**x**_0_) ((S18)) from avian gastrulation experimental data discussed in main text Fig. 4, with Λ(**x**_0_) taken as the largest of the forward and backward time FTLEs, corresponding to the smallest Lyapunov time (See Formula in Algorithm 1).

**Fig. S5:**
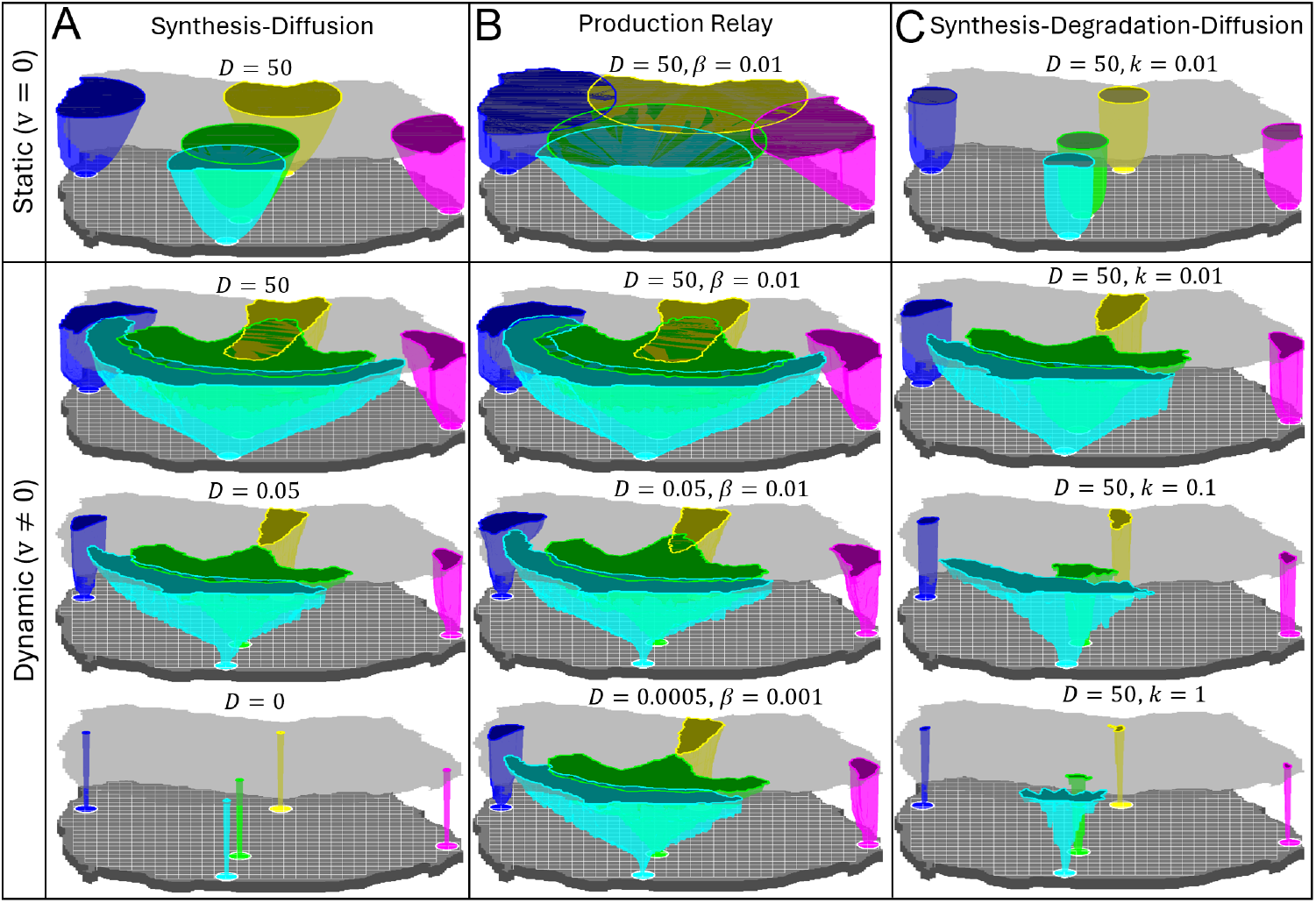
Embryological Light Cones in Avian Gastrulation. Same as ELCs in main text Fig. 4 with alternative signaling mechanisms and parameters in static (top row, **v** = 0) and dynamic (bottom rows) using chick gastrulation velocities. Sources are as in main text Fig. 4. Final contour is shown at *T* = 12*h*. All parameters are in *µm* and *min*. Concentration units are arbitrary with a secretion rate of 100*min*^−1^. A) Synthesis Diffusion (as in the main text Fig. 4), illustrating sub-linear static expansion and reduced ranges as diffusion decreases. B) Production relay, as in A, but with the addition of a logistic growth term (+*βc*(*c*_max_ − *c*), *c*_max_ = 100). ELCs expand linearly with positive feedback, increasing interaction ranges, even at low diffusivity. C) SDD (Synthesis-Degradation-Diffusion) model, as described in the main text Fig. 2, but with moving point sources (as in A) and a secretion rate replacing the fixed flux boundary condition. ELCs approach a fixed range in the static case (top) but can continue expanding in the dynamic case due to convergent tissue deformation. In all cases, ELCs are augmented by deformation in robust, predictable ways as long as *D* > 0. Overall, dynamic tissues affect cell-cell interaction ranges, with all of the important qualitative features of ELCs’ distortion predictable from **C**.

### Movies

Movie 1: Time evolution associated with Fig. 1C.

Movie 2: Time evolution associated with Fig. 2E-H.

Movie 3: Time evolution associated with Fig. 2I-K.

Movie 4: Deformation of randomly colored tissue patches in avian gastrulation. Same velocity data as Fig. 4.

Movie 5: Time evolution associated with Fig. 4A-G.

